# Social ‘envirotyping’ the ABCD study contextualizes dissociable brain organization and diverging outcomes

**DOI:** 10.1101/2024.08.20.608873

**Authors:** Haily Merritt, Mary Kate Koch, Youngheun Jo, Evgeny J. Chumin, Richard F. Betzel

## Abstract

The environment, especially social features, plays a key role in shaping the development of the brain, notably during adolescence. To better understand variation in brain-environment coupling and its associated outcomes, we identified “envirotypes,” or different patterns of social environment experience, in the Adolescent Brain Cognitive Development Study by hierarchically clustering subjects. Two focal clusters, which accounted for 89.3% of all participants, differed significantly on eight out of nine youth-report social environment quality measures, representing almost perfect complements. We then applied tools from network neuroscience to show different envirotypes are associated with different patterns of whole brain functional connectivity. Differences were distributed across the brain but were especially prominent in Default and Somatomotor Hand systems for these focal clusters. Finally, we examined how social envirotypes change over development and how these patterns of change are associated with a suite of outcomes. The resulting dynamic envirotypes differed along dimensions of stability and quality, but outcomes diverged based on stability. Specifically, the stable, high quality envirotype was most easily distinguished from the improving envirotype, while the unstable envirotype was associated with the worst outcomes. Altogether, our findings represent significant contributions to both social developmental neuroscience and network neuroscience, emphasizing the variability and dynamicity of brain-environment coupling and its consequences.

## INTRODUCTION

More than an arena for social cognition to play out, the social environment is home to resources important for human cognition and and wellbeing [17, 22, 36, 52, 97]. The quality of social relationships and the support received from them has a considerable impact on health [4, 20], including reduced physiological impact of adversity [93], better prognoses related to disease [18– 20, 92], increased antibody response to vaccines [85], lower levels of inflammation [86], conserved energy in response to threats [33, 73], decreased risk of all-cause mortality [5, 89], and more [16, 38, 44, 50, 69, 75, 84]. Indeed, many facets of the social environment have overlapping influences [65]. More than that, social support in particular can buffer against the otherwise harmful effects of adverse (social) environmental experiences [41, 52, 64, 70, 73, 94]. Such findings emphasize the pervasive impact of the social world on every facet of human neuroscience, physiology, cognition, and behavior.

Accordingly, neuroscientists have consistently identified differences in both mental health and cognitive outcomes associated with different brain network organizations in different social environments [26, 64, 65, 67]. Indeed, brain network differences have been found for differences in childhood abuse and neglect [45, 63, 81], childhood maltreatment [24], institutional care [40], occurrence of stressful or traumatic life events [56], exposure to violence [81], educational attainment [15, 65, 68], occupational prestige [15], household poverty status or income [26, 65, 78], neighborhood or area disadvantage or deprivation [64, 65, 68], positive social support [52, 60, 94], family or peer adversity [72, 88], and loneliness [53, 54, 77], among others [39, 42]. Perhaps most notable across these findings is that the particular pattern of brain network organization associated with better mental health and better cognitive performance *depends on the environment* [26, 67]. The contextuality of brain network organization and associated outcomes is a feature, not a bug, of human neuroscience.

Beyond varying across contexts, brain networks reconfigure over the course of development [6, 59] in accordance with experience in the environment [1, 9, 26, 43, 67]. Several studies have suggested that much of the observed variation in brain network organization across individuals can be linked directly to variation in environmental experience[1, 15, 31, 67, 95], where brain network reconfiguration over development shapes and is shaped by the environment [1, 13]. In other words, certain patterns of brain network organization are consistently associated with certain trends in environmental experiences, especially during adolescence [8, 29, 71]. For clarity and consistency, we refer to a particular pattern of environmental experiences here as an envirotype. For example, individuals from social envirotypes characterized by less social support had denser connectivity between Default and Visual, Somatomotor, and Ventral Attention Networks than their more supported peers [53]. Brain network organization calibrates in alignment with an envirotype over time to support adaptive behavior and physiological responses, meaning individuals in different environments will engage behaviorally, neurally, and physiologically with their environments differently [1, 9, 23, 48, 67]. A critical consequences of this is that individuals whose brain network organization diverges from the norm *for their envirotype*—as opposed diverging from a “universal” norm—are at greater risk for poorer mental health outcomes [67]. Contextualizing brain network differences in this way is one route to reconcile incompatibilities in brain network-based biomarkers of, for example, depression [10, 61]. Put another way, by studying envirotypes and their associations with different patterns of brain network organization and diverging outcomes, we develop a richer and more nuanced understanding of brain-environment coupling which helps researchers and policy-makers better serve at-risk youth [90].

Rigorously quantifying the dynamics of envirotype-brain network associations requires large longitudinal samples, which can be prohibitively expensive in terms of both money and time. Accordingly, many researchers have taken advantage of the Adolescent Brain Cognitive Development (ABCD) study [2, 14, 30], especially elucidating the association between socioeconomic status (SES) and brain network organization [26, 64, 65]. Since the social environment has such a pervasive influence on development during adolescence [29], it is important to consider a diversity of social environmental measures beyond distal, structural measures like SES. Because subjective social support—specifically, perceived social support’s alignment with expectations and desires for support—provides buffering effects against socioeconomic disadvantage [11, 52, 73, 94] and can provide greater predictive power than more objective SES measures [17, 21], this facet of the social environment is especially critical for neuroscientific investigation.

In line with this goal, we leverage the size and longitudinality of the ABCD dataset to examine associations between social envirotypes and brain network organization over adolescence using a suite of measures, as well as the behavioral, cognitive, life experience, and clinical outcomes of these associations. We use data-driven techniques to cluster thousands of adolescents at baseline using their social environmental experiences, resulting in social envirotypes. We then apply tools from network neuroscience to clarify how patterns of whole brain functional connectivity differ across these baseline envirotypes. Finally, we quantify the change in social envirotypes over several years during the critical transition period of adolescence using longitudinal social environment data. We show how different patterns of change in the social environment are associated with different outcomes for mental health, cognition, behavior, and life experiences. By contextualizing brain network organization and outcomes in terms of dynamic social envirotypes, these findings contribute to the nuanced narrative of brain-environment coupling and its consequences.

## RESULTS

Our **Results** section is organized as follows. First, we describe a data-driven procedure for identifying envirotypes at baseline in the ABCD study. We then show that these envirotypes exhibit differences in brain network organization as indexed by functional connectivity. Finally, we identify the different dynamics of envirotypes over development using longitudinal social environment measures, as well as their diverging outcomes on a suite of clinical, cognitive, behavioral, and life experience measures.

### A. Envirotypes represent qualitatively different social environments

Previous studies have investigated the association between social environment and brain network organization and outcomes using the ABCD dataset, often focusing on socioeconomic status because of its ease of measurement [26, 65]. Such work has tended to define groups according to high or low scores on one or two measures, such as parent income or education. While this approach has been remarkably informative, many questions remain. Here, we leverage the size of the ABCD dataset to better approximate the high dimensionality of the social environment and the variability therein to extract “envirotypes.” These envirotypes represent distinct patterns of social environment experience, which we then link to different patterns of brain organization and diverging outcomes. We opt for *subjective* measures of social environment *quality*, since recent work suggests they may have stronger associations with neural data [17, 21].

We first identified baseline envirotypes using all youth-report measures of social environment quality with at least an 80% response rate at the first time point of data collection among subjects with high-quality neuroimaging data (*N* = 2776 subjects; see Figure S1 for a comparison of social environment scores for those with and without quality imaging data). Through this selection process, we arrived at nine measures reflecting family, school, and peer relationships: Community Risk, Family Conflict, School Involvement, Neighborhood Safety, Caregiver Acceptance, School Disengagement, Parental Monitoring, School Environment, and Social Resilience (see Figure 1a for subjects’ scores, Figure S2 for measures’ correlations with each other, and Supplementary Table 1 for measure means). We computed the pairwise correlations between all subjects across these nine measures (see Figure 1b), and applied a hierarchical clustering algorithm to the resulting correlation matrix [7, 62]. The hierarchical nature of this clustering algorithm allows us to change the resolution of the clusters, such that lower hierarchical levels correspond to broader, more heterogeneous clusters and higher hierarchical levels correspond to increasingly narrow and homogeneous clusters.

**Figure 1.**
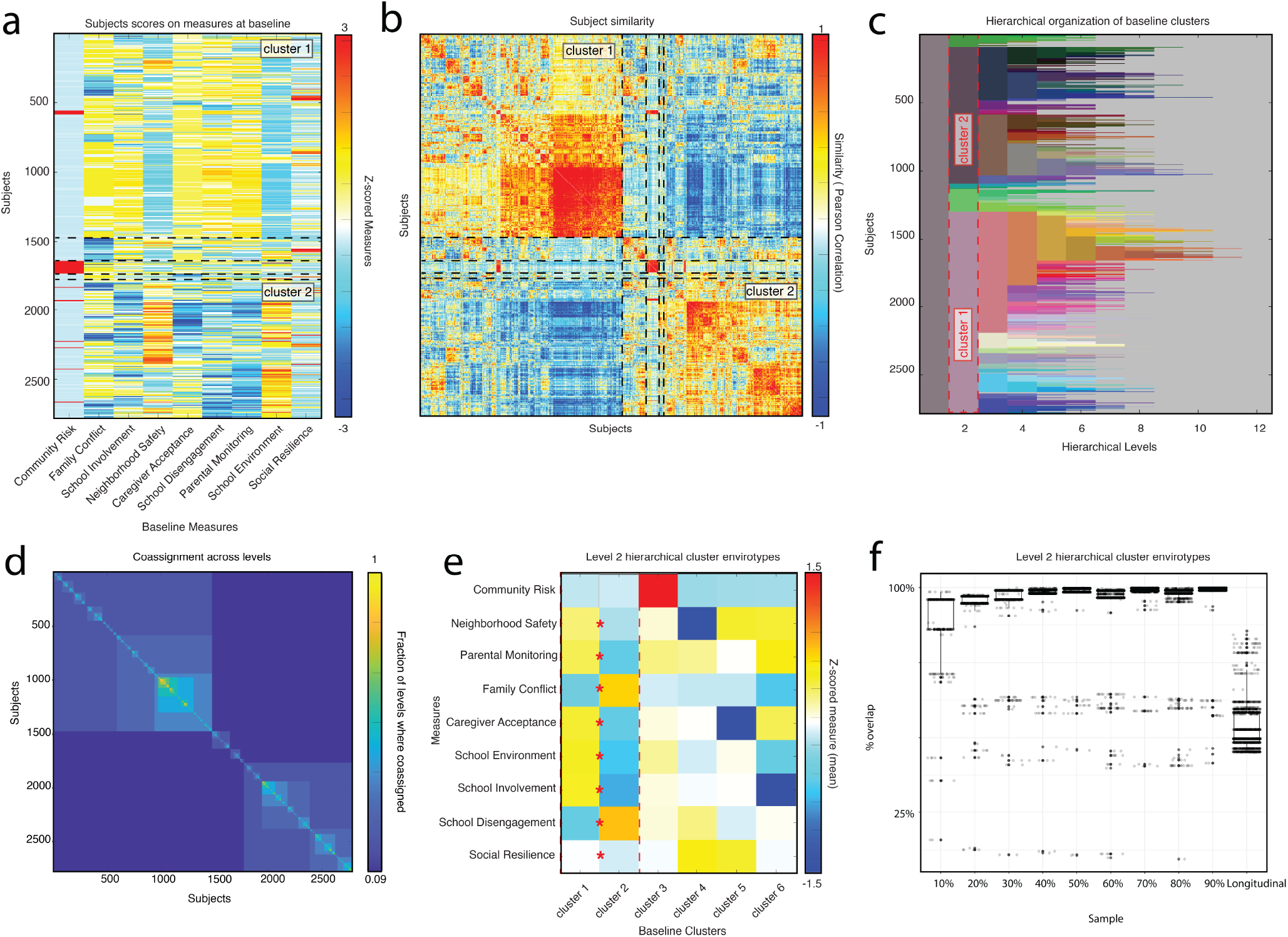
Hierarchical envirotypic clusters at baseline. Panel (a) shows all subjects scores across nine social environment quality measures recorded at the first time point of data collection, ordered by level 2 baseline cluster membership. Red indicates a higher quality environment for all measures except Family Conflict and School Disengagement. Panel (b) shows pairwise similarity of all subjects to each other using Pearson correlation, ordered by level 2 baseline cluster membership. Panel (c) shows subjects’ baseline cluster membership across all hierarchical levels. Panel (d) shows the number of hierarchical levels in which subjects were assigned to the same cluster, ordered by level 2 baseline cluster membership. Panel (e) shows cluster-averaged Z-scores on nine social environment quality measures for the six clusters present in level 2. Panel (f) compares the percent overlap in people assigned to the same cluster across a range of subsamples, as well as the subjects included in the longitudinal clusters (Fig. 3) to assess the validity of our identified baseline clusters. With the exception of the longitudinal sample, all subsamples had a mean percent overlap *>* 0.875, suggesting our identified baseline clusters were reasonable representatives of the envirotypes described. The mean percent overlap for the longitudinal sample was 0.533 *±* 0.081, which we expect to be lower since the longitudinal clusters capture envirotype dynamics.

Our clustering algorithm yielded 12 hierarchical levels, representing different resolutions of baseline clusters (see Figure1c and d). We focus primarily on hierarchical level 2 (but see Supplementary Figure S5 for cluster refinement across levels and Supplementary FigureS6 for envirotypic comparison across levels), which contained six clusters comprised of 1475, 1005, 91, 165, 36, and 4 participants each (representing 53.1, 36.2, 3.3, 6, 1, and 0.1% of the total sample; see Figure1c). These clusters differed in their composition across hierarchical levels, such that Baseline Cluster 1 specialized into an increasingly narrow envirotype while Baseline Cluster 2 immediately fractures into distinct clusters at a finer resolution (see Supplementary Figure S6 for envirotypes at finer resolutions and Supplementary Figure S7 for brain network differences between finer resolution envirotypes).

We furthermore focus on these two largest clusters at hierarchical level 2, which collectively accounted for 89.3% of all participants. Baseline Cluster 1 presented an envirotype characterized by especially high Parental Monitoring, low School Environment, and low Neighborhood Safety. The envirotype of Baseline Cluster 2 was characterized by especially high Neighborhood Safety, high School Environment, and low Parental Monitoring (see Figure1a,e). These two largest clusters were significantly different on all measures *except* Community Risk, representing near-perfect complements of each other (all *p*s *<* 0.04; *t*-tests corrected for multiple comparisons using the Benjamini-Hochberg method, accepted FDR = 0.05, adjusted *p*_*crit*_ = 0.046). These clusters were relatively stable across a range of subsampling (see Figure 1f for subsample percent overlap of community membership and Supplementary Figure S4 for centroids of subsampled clusters).

One possibility is that the detected envirotypes simply reflect other well-studied but indirect measures of environment, such as SES. To rule this out as a possible interpretation, we examined the distribution of SES across the baseline clusters. As a measure of SES, we used the first principal component of a principal component analysis applied to 14 measures of SES, including parent income and parent education (see Supplementary Table S2 for measures and their descriptive statistics). This first principal component explained 56.5% of the variance, whereas the second component explained only 13.1% of the variance (see Supp. Fig. S3f). An ANOVA indicated significant pairwise differences, but using a post-hoc Tukey’s test we found no significant difference in composite SES between focal clusters 1 and 2 (*p* = 0.362). We did find, however, a significant difference in composite SES between clusters 1 and 4 (*p* = 0.007).

We also investigated whether pubertal status or age was associated with baseline cluster composition, but there were no differences in either pubertal status or age between any pairs of clusters using a one-way ANOVA. Additionally, we tested the correlations of these youth-report measures with the corresponding caregiver-report measures where available. All three were positively correlated, though the magnitude varied (*r* = 0.88, *p* = 2.2*e −* 16 for Community Risk; *r* = 0.31, *p* = 2.2*e −* 16 for Neighborhood Safety; *r* = 0.20, *p* = 2.2*e −* 16 for Family Conflict).

Given that our clustering algorithm is hierarchical, we also examined finer resolution clusters (i.e., the red and green clusters that appear on level 3 in Figure 1c). These clusters were characterized by diverging experiences in family versus school domains, where one envirotype had a relatively more negative family but relatively more positive school experience and vice versa for the other envirotype (see Supplementary Figure S6 and Supplementary Figure S7a). This result suggests different domains of the social environment can have different influences, in accordance with Bronfenbrenner [12].

In summary, we identified dissociable envirotypes based on measures of social environment quality at baseline. These envirotypes reflect different patterns of perception of social environment experience during early adolescence. In addition, they may align with other constructs in youths’ developmental contexts, such as the control and support dimensions of parenting styles [46]. Baseline Cluster 1 displayed an envirotype characterized by higher Parental Monitoring but less Family Conflict, less School Disengagement and a supportive School Environment, and high Neighborhood Safety. Baseline Cluster 2’s envirotype was defined by less Parental Monitoring and greater Family Conflict, lower quality School Environment and more School Disengagement, and less Social Resilience and Neighborhood Safety. Moreover, the envirotypes we identified capture features of the environment not fully explainable by socioeconomic status, age, pubertal development, or data collection site.

### B. Brain network organization differs between envirotypic clusters

Next we sought to characterize how the focal envirotypes differed in their brain network organization at the same time point (see Figure 2a and b for representations of the brain network organization of each corresponding cluster, Figure 2c for their difference, and Supplementary Figure S8a and b for the standard deviation of edge weights for each cluster). To do this, we first performed mass *t*-tests on each edge weight, correcting for multiple comparisons using the Benjamini-Hochberg method (accepted FDR = 0.05, adjusted *p*_*crit*_ = 0.04. These tests yielded significant differences between clusters for much of the Somatomotor Hand system, as well as connectivity within Auditory, Cingulo-Opercular, and Default systems (see Figure 2d).

**Figure 2.**
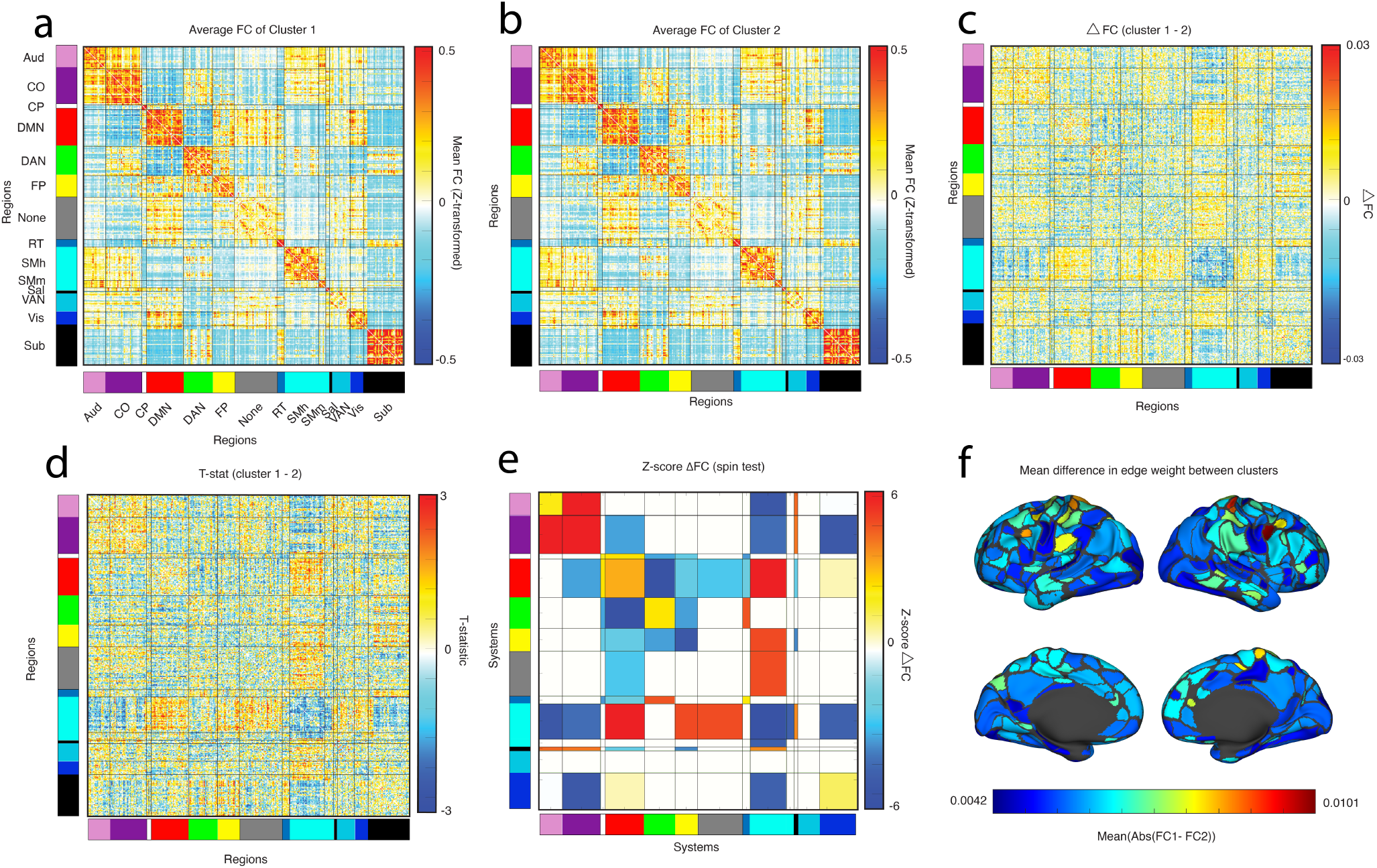
Differences in functional connectivity (FC) between clusters. Panel (a) shows average FC for all members of cluster 1 in level 2. Panel (b) shows average FC for all members of cluster 2 in level 2. Panel (c) shows the difference between clusters 1 and 2’s FC (i.e., the difference between panel a and b). Panel (d) shows the T-statistic for each edge of the difference between clusters 1 and 2. Panel (e) the system-level summaries of a ‘spin test’ between the FC of clusters 1 and 2. Panel (f) shows the nodal mean of the mean squared difference in edge weights between clusters. In other words, brighter red indicates that a node’s connections tended to have large magnitude differences between clusters, whereas darker blue indicates negligible differences in connectivity between clusters.

We wanted to test whether there were system-level effects, so we calculate the mean difference in FC for all system-by-system blocks. We compare each blockwise value against a null distribution generated using cortical spin tests, which preserve spatial statistics. FC differences were statistically significant if the observed difference exceeded that of the null distribution (false discovery rate fixed at *q* = 0.05; adjusted critical value of *p*_*crit*_ = 0.02). The “spin” test showed that Baseline Cluster 1 (characterized by an envirotype with higher Parental Monitoring) had greater connectivity especially within Cingulo-Opercular and Default systems than did Baseline Cluster 2. Baseline Cluster 1 also had greater connectivity between Cingulo-Opercular and Auditory systems and between Somatomotor Hand and Default, Somatomotor Hand and Frontoparietal, Somatomotor Hand and Salience, Salience and Auditory, Salience and Cingulo-Opercular, and Retrosplenial-Temporal and Dorsal Attention. Conversely, Baseline Cluster 2 (characterized by an envirotype with higher Family Conflict) had greater connectivity within Frontoparietal and Somatomotor Hand. Additionally, Baseline Cluster 2 had greater connectivity between Default and Dorsal Attention, Somatomotor Hand and Auditory, Somatomotor Hand and Cingulo-Opercular, Somatomotor Hand and Visual, and Visual and Cingulo-Opercular (all *p*s *<* 0.02; see Figure 2e). Put succinctly, Baseline Cluster 2 largely had stronger FC between somatosensory systems and within Somatomotor Hand. Many of the largest differences between clusters were found in the connectivity profiles of the Somatomotor Hand and Default systems.

To further understand the distribution of differences in FC between envirotypic clusters, we computed the mean squared difference between the clusters for each edge weight. This procedure yields a matrix whose dimensions are the number of nodes by the number of nodes, where each element is the mean squared difference. We take the mean of each row of this matrix, which gives a single value for each node, and plot in anatomical space to identify where in the brain large differences between clusters are (see Figure 2f and Supplementary Figure S8c-f for different calculations of nodal differences plotted in brain space). Again, we see that the two focal clusters are distinguished by patterns of connectivity involving Default and Somatomotor regions, while Visual connectivity is strongly preserved.

Additionally, we examined differences in brain network organization between finer resolution clusters. As with the level 2 baseline clusters, we performed mass t-tests on each edge weight, correcting for multiple comparisons using a false discovery rate of 0.05, and a space-preserving spin test. Again, we found differences distributed across the brain. Worse family environment was associated with higher connectivity between Auditory and Visual, CO and DMN, Somatomotor and DMN, and within VAN. Worse school environment was associated with increased connectivity between Auditory and CO, within DMN, CO and VAN, VAN and Visual, and Salience and DAN (see Supplementary Figure S7).

In summary, differences in brain network organization between envirotypes are distributed widely across the brain at baseline. Coarser envirotypes, which have a clearer positive versus negative distinction, differ strongly in Somatomotor and Default connectivity. Finer envirotypes, which differ more in which domain carries which valence, display differences in VAN and CO connectivity.

### C. Envirotypic clusters are dynamic and associated with a suite of outcome variables

Because adolescence is a period characterized by rapid biological and social change, we sought to understand how social envirotypes change over time and the impact this change has on behavioral, cognitive, clinical, and life experience outcomes. To do this, we applied our hierarchical clustering algorithm to longitudinal measures of social environment quality. We used six measures of social environment quality, a subset of our previously clustered measures, which represented all measures of social environment quality with at least an 80% response rate across all time points (see Figure 3a). We proceeded with the same methods as before, calculating the correlation between all subjects across the 18 scores (i.e., six measures at three time points; see Figure 3b) and the applying our hierarchical clustering algorithm to the resulting matrix of similarities. Our hierarchical and longitudinal clustering yielded 11 hierarchical layers, representing different levels of resolution (see Figure 3c and Supplementary Figure S9 for cluster refinement across levels). We focus primarily on level 2, which had six clusters, with 614, 301, 382, 223, and 7 subjects, respectively (see Supplementary Figures S10, S11, S12,and S13 for finer resolution data for each of the larger four clusters). We excluded Longitudinal Cluster 5 from the subsequent analyses because it was too small for statistical comparisons.

**Figure 3.**
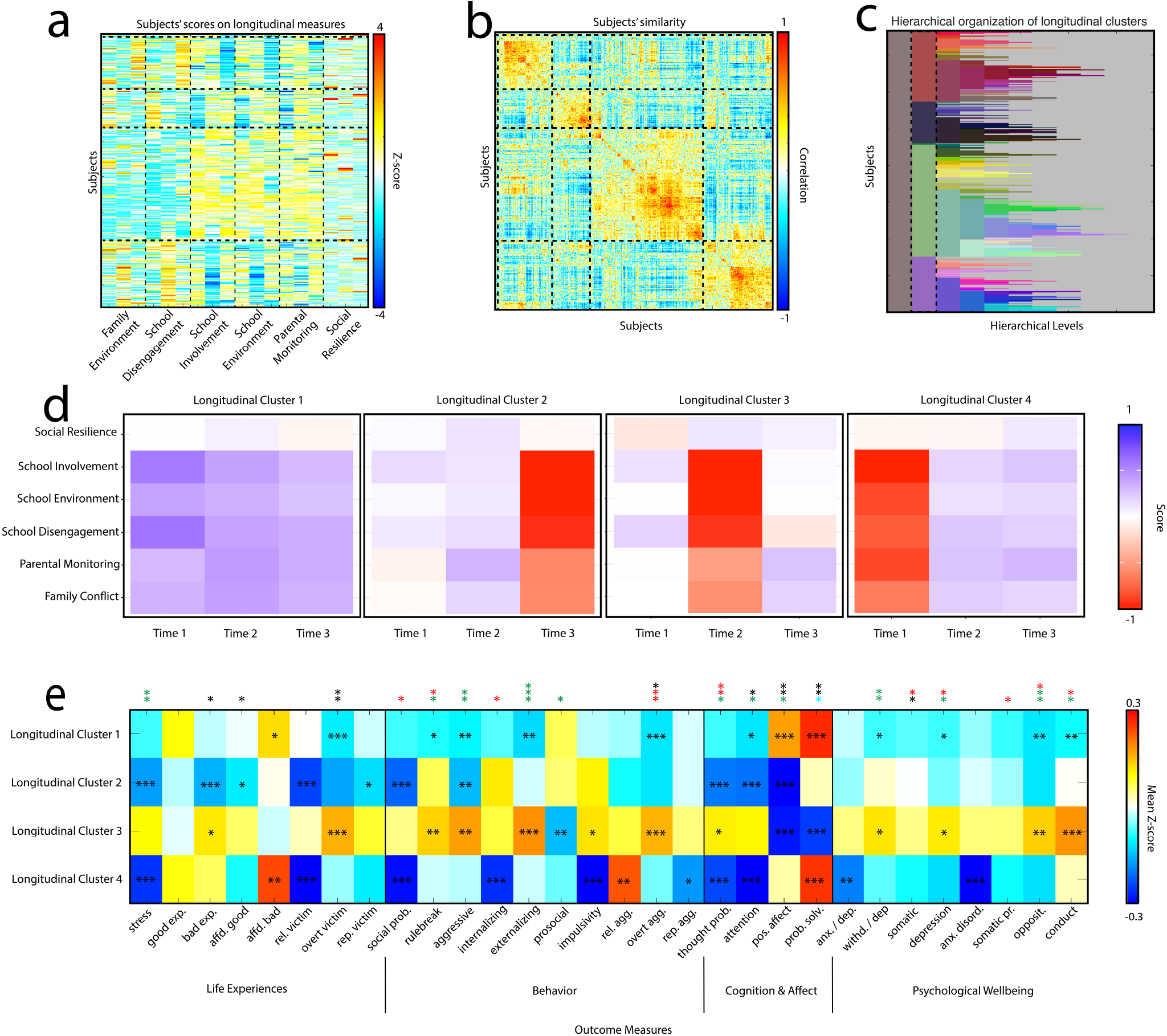
Social envirotypes over time. Panel (a) shows subjects’ z-scores on six measures of social environment quality over three time points during adolescence. Horizontal dashed lines separate clusters, while vertical dashed lines separate measures. Panel (b) is the similarity matrix, achieved by calculating the pairwise Pearson correlation between all subjects across the six measures of social environment quality at three time points. Dashed lines of both orientations separate clusters. Panel (c) shows the cluster assignments across hierarchical levels for the longitudinal data. Cluster assignments in other panels refer to level 2, which is outlined with a dashed box. Panel (d) shows the envirotype dynamics of each cluster. Note that Longitudinal Cluster 5 is excluded because it has too few members for statistical comparisons. Panel (e) shows how each of the longitudinal envirotypes score on 30 measures of life experience, behavior, cognitive/affective, and psychological wellbeing outcomes. Asterisks within cells indicate that a given cluster has a more extreme score than would be expected by chance, determined by a permutation test with 1000 permutations. Asterisks on top indicate significant comparisons between clusters, determined by a one-way ANOVA and post-hoc Tukey’s test. Black asterisks represent comparisons between Longitudinal Clusters 1 and 4; red asterisks represent comparisons both between 1 and 4 and between 2 and 4; green asterisks represent comparisons between 1 and 4, 2 and 4, and 3 and 4; and blue asterisks represent comparisons between 1 and 2, 1 and 4, 2 and 4, and 3 and 4. *: *p <* 0.05, **: *p <* 0.01, ***: *p <* 0.001.

The envirotypes associated with the identified longitudinal clusters were generally distinguished by the stability and perceived quality of the social environment, broadly in line with previous theoretical work in this space [76, 83]. Longitudinal Cluster 1 was characterized by a stable, positive social envirotype, consistently scoring high on measures of positive social environment quality (e.g. School Involvement, School Environment, and Parental Monitoring) and low on measures of negative social environment quality (e.g. Family Conflict and School Disengagement; see Figure 3d). All other clusters displayed instability in their envirotypes. Longitudinal Cluster 2 worsened over time, while Longitudinal Cluster 4 improved over time. Longitudinal Cluster 3 was more generally unstable, with a relatively negative social environment at the second time point (see Figure 3d).

Finally, we examined how these dynamic envirotypes are associated with a suite of outcomes, using all cognitive, clinical, behavioral, and life experience measures with at least an 80% response rate within our sample at the last time point, resulting in a total of 30 outcomes (see Figure 3e and Supplementary Figure S2 for associations with social environment quality measures). We first performed a one-way ANOVA for each outcome to identify differences between clusters. For significant comparisons, we performed a posthoc Tukey’s test, correcting for multiple comparisons using a false discovery rate of 0.05. All significant differences involved at least Longitudinal Clusters 1 and 4, which were characterized by Stable and Improving envirotypes, respectively. These envirotypes diverged on 21 out of 30 outcomes, with the stable envirotype seeing better outcomes on all measures except Social Problems, Internalizing, Thought Problems, and Depression. Longitudinal Clusters 2 and 4—Worsening and Improving Envirotypes, respectively—differed on 18 out of 30 measures, with the worsening envirotype seeing better outcomes on all measures except Rulebreaking, Internalizing, Positive Affect, Problem Solving, and Withdrawal/Depression. Longitudinal Clusters 3 and 4—Unstable and Improving envirotypes, respectively—differed on 12 out of 30 measures, with the improving envirotype having better outcomes on all measures except Attention (see Figure 3e). In general, although the clearest divergence in outcomes was between the stable and improving envirotypes, the worst outcomes were consistently associated with the unstable envirotype.

To further understand the outcomes associated with the four different dynamic envirotypes, we performed permutation testing to characterize the extent to which a given cluster was saturated with extreme scores. Longitudinal Cluster 3 (Unstable Envirotype) was especially saturated with worse outcomes on 15 out of 30 measures. The cognition and affect measures clearly distinguished between clusters, with all clusters being especially saturated for 3 out of 4 of this subset of measures (see Figure 3e).

In summary, dynamic envirotypes were distinguished by both the quality and stability of the social environment over the course of adolescence. Stable and Improving Envirotypes were most clearly different from one other in terms of outcomes, with the Stable Envirotype tending to have better outcomes. The Unstable Envirotype, however, consistently had the worst outcomes.

## DISCUSSION

We use social envirotypes to contextualize differences in brain network organization and diverging outcomes across a suite of measures. We combine a framing motivated by developmental and social neuroscience with data-driven analyses to rigorously quantify and characterize these hierarchical and dynamic envirotypes. Specifically, we used measures of perceived social environment quality to cluster adolescent subjects from the ABCD dataset at baseline. These baseline clusters captured envirotypes (i.e., patterns of social environment experience), with subjects from focal clusters differing on 8 out of 9 measures. We found differences in brain network edge weights between the two focal clusters. These differences were distributed across the brain, but the strongest and most pervasive effects involved Somatomotor and Default systems. Additionally, we leveraged longitudinal data from ABCD to examine how envirotypes changed across adolescence, linking this change to outcomes of psychopathology, behavior, cognition, and life experiences. We focused on four longitudinal clusters, whose envirotypes were characterized by stable high quality, general instability, improvement, and worsening. The outcome measures most clearly differentiated the stable and worsening envirotypes, while the unstable envirotype exhibiting the worst outcomes. Altogether, these results depict a contextual and dynamic story about the coupling between brain networks and the social environment. By studying envirotypes and their associations with different patterns of brain network organization and diverging outcomes, we contribute to a richer and more nuanced understanding of brain-environment coupling.

### D. Envirotypes are a way to contextualize variation

By starting our investigation from the envirotypes— which are themselves defined by a suite of social environment quality measures—we shift the emphasis from individual differences (i.e., people are different from one another) to contextualizing difference and variation (i.e., people are different from one another in part because their environments are different). Broadening the scope of study in this way facilitates clearer understanding of variation, its potential influences, and its consequences. By contextualizing variation, we can identify not only who is at risk of, for example, depression but also how to more effectively intervene on the context to ameliorate such a risk [67]. Put another way, by emphasizing the context of phenotypes (e.g. brain network organization) and their outcomes we can target interventions at the environment in addition to the individual. We note that while we may be the first to use the term ‘envirotype’ in the context of developmental, social, and network neuroscience, it has previously been used in machine learning contexts linking genotypes, phenotypes, and envirotypes[37].

We first identified envirotypes based on patterns of social environment experience using a hierarchical clustering algorithm. By identifying clusters hierarchically, we can circumvent having to choose a single resolution for study a priori [28]. We focus on two baseline clusters at a coarse level whose envirotypes were complementary: safer, more resourced, and more supportive on the one hand and less safe, less accepting, and involving more conflict on the other. It is possible these clusters partially reflect differences on the control and support dimensions of parenting styles, with adolescents in cluster 1 tending to have parents higher on support and control (i.e., authoritative style) and adolescents in Baseline Cluster 2 tending to have parents lower on support but higher on control (i.e., authoritarian; 3, 46). Differences in parenting styles may also have a bidirectional influence with environment quality, as prior work suggests negative effects of high controlling parenting styles contextually vary (e.g., in high-risk neighborhoods; 79). We note that, while at a coarse level this distinction between more and less “socially desirable” envirotypes is clear, at finer resolutions the distinction is blurred. This increasing nuance with finer resolutions speaks to the importance of resolution-agnostic methods, like hierarchical clustering [7].

### E. Brain network organization differs between social envirotypes

With two distinct envirotypes defined via data-driven clustering, we address how brain networks are organized differently for different envirotypes. We find differences distributed widely across the brain, yet nearly 40% of all system pairs with significant differences involved either the Default or Somatomotor Hand systems. Generally, Baseline Cluster 2—which was characterized in part by lower Neighborhood Safety and Caregiver Acceptance– had more positive connectivity between somatosensory systems and within the Somatomotor Hand system than Baseline Cluster 1. Importantly, brain systems whose connectivity profiles differed based on social environment quality were not limited to those systems implicated in social cognition (e.g., Default, Salience). Indeed, *both* somatosensory systems and systems associated with higher order cognition were dissociable, in line with several previous studies [64–66]. While much previous work has focused on environment-associated connectivity differences in regions associated with higher order cognition [26], differences in somatosensory regions are consistently identified between groups from different social environments [53, 65]. While theoretical work on trauma and neurodevelopment provides explanations as to why, for example, amygdala and prefrontal connectivity should vary with social environmental experience [83], it is unclear how variation in somatosensory connectivity in accordance with the social environment may be adaptive. Nevertheless, our findings reiterate several others’ results: brain-environment coupling involves the whole brain. This refrain motivates a move away from studying connectivity between only a few regions of interest and toward taking advantage of the methodological tools from, for example, network science [49, 57]. Beyond suggesting methodological approaches, these findings on brain network differences underscore that although some variation in brain network organization can be explained by disease status or genetics [8, 71], calibration to the environment represents a critical source of variation in brain network organization [1, 55].

A reason we see different patterns of connectivity as contingent upon different social experiences might be that they support different behaviors and neurophysiological responses that are adaptive in their respective environments [26, 68]. This perspective echoes arguments made about the centrality of experiencedependent plasticity in the development of the nervous and stress response systems [9, 23, 25]. Experiencedependent plasticity implies there are many possible neuroendophenotypes of brain network organization. Which one manifests for a given individual depends on experience. Moreover, what is an adaptive pattern of network organization in one context may not be adaptive in another context, or even in the same context at another stage of development. Given this, we might expect network organization to mediate the association between experience in the environment and mental health outcomes, as demonstrated by previous work [26, 56, 68]. Put another way, phenotypic divergence from the norm *for one’s environment* may better predict risk than divergence from a global norm, since different behaviors are better suited for different environments [67].

Moreover, by examining the context of brain network differences and diverging outcomes, we open up new possibilities for intervention to improve wellbeing and rectify harm. Indeed, recent work suggests “difficult lives explain depression better than broken brains” [51]. This argument is compounded by studies showing that the patterns of brain organization associated with depression and other outcomes depend on the environment [26, 67], which clarifies work proposing conflicting brain network-based biomarkers of, for example, depression [10, 61]. Put more simply, there may be no one brain network-based biomarker for psychopathology. With this in mind, contextualizing brain network organization seems to be a critical area for future network neuroscience research, especially investigations of resilience to stress [47]. Here we emphasize context in terms of social environment, but there are many possible avenues for contextualization. Work by Tian and colleagues casts different bodily systems as the context to understand aging and common neuropsychiatric disorders [82, 91]. Recent work outlining the differing paces of development across the brain suggests considering time as context is important as well [80].

### F. Different envirotype trajectories have diverging outcomes

Finally, we leveraged the longitudinality of the ABCD dataset to determine how envirotypes change over time and what kinds of outcomes are associated with these patterns of change. Our clustering algorithm suggested four general patterns of change: high quality stability, instability, improvement, and worsening. That is, not only the quality but also the perceived stability of that quality is important for differentiating envirotypes and their dynamics. This data-driven result is in accordance with theoretically oriented research in developmental neuroscience, which suggests that dimensions of harshness and instability are especially powerful influences on the developing brain and later wellbeing [76, 83]. While the Stable and Improving Envirotypes had the most distinct outcomes—with the Stable Envirotype generally but not exclusively seeing more positive outcomes—the Unstable Envirotype was consistently associated with the worst outcomes. These findings suggests that it not only matters how we perceive where we start, but also how robust that perception is over time. Additionally, these results point to the importance of the timing of a measurement when associating it with outcomes, especially since environmental instability is notably difficult to measure effectively [96].

Focusing on variation—whether of phenotypes, outcomes, or trajectories—supports the neural, psychological, and behavioral sciences in their goal of better understanding humans [22, 58, 74]. Ignoring the dynamic environmental context in which humans are situated delays not just scientific clarity but also effective interventions for wellbeing. Just as we are in our environments, our environments are “in” us.

### G. Limitations and Future Directions

Although ABCD is perhaps the largest and most diverse publicly available sample of adolescents, the extent to which our results represent global patterns of envirotypes and envirotypic change is unclear since all subjects are US-based. Additionally, because our neuroimaging data are from one time point, we cannot speak to how envirotypes, dynamic or not, are associated with brain network reconfiguration across development. Association-based approaches like ours are also unable to comment on the direction of causality between envirotypes and brain network variation. Future research can answer this critical question by leveraging additional time points of neuroimaging data from ABCD as they are released. While we used a suite of social environment quality measures to capture the richness and nuance of the social environment, alternative measures may identify different patterns of brain-environment coupling or envirotype dynamics.

## H. Conclusions

Using a data-driven approach, we identified baseline envirotypes that differed across eight measures of perceived social environment quality. These baseline envirotypes displayed dissociable patterns of brain network organization, largely featuring somatosensory and default regions. When we examined how envirotypes change over time, we found four predominant patterns: stable high quality, unstable, improving, and worsening. The stable and improving envirotypes had the greatest differences on a suite of behavioral, cognitive, clinical, and life experience outcomes, while the unstable envirotype showed the displayed the worst outcomes. Altogether, our findings represent significant contributions to both social developmental neuroscience and network neuroscience, emphasizing the variability and dynamicity of brain-environment coupling and its consequences.

## I. Acknowledgements

Data used in the preparation of this article were obtained from the Adolescent Brain Cognitive DevelopmentSM (ABCD) Study (https://abcdstudy.org), held in the NIMH Data Archive (NDA). This is a multisite, longitudinal study designed to recruit more than 10,000 children age 9-10 and follow them over 10 years into early adulthood. The ABCD Study® is supported by the National Institutes of Health and additional federal partners under award numbers U01DA041048, U01DA050989, U01DA051016, U01DA041022, U01DA050987, U01DA041117, U01DA050988, U01DA041025, U01DA041148, U01DA051018, U01DA041174, U01DA041028, U01DA051039, U01DA041120, U01DA041093, U01DA051037, U01DA041106, U01DA041134, U01DA041156, U01DA051038, U01DA041089, U24DA041123, U24DA041147. A full list of supporters is available at https://abcdstudy.org/federalpartners.html. A listing of participating sites and a complete listing of the study investigators can be found at https://abcdstudy.org/consortium_members/. ABCD consortium investigators designed and implemented the study and/or provided data but did not necessarily participate in the analysis or writing of this report. This manuscript reflects the views of the authors and may not reflect the opinions or views of the NIH or ABCD consortium investigators.

## METHODS

### J. Social environment data

We used social environment quality and resting state fMRI data from 2776 (1418 female) adolescents age 8.9 - 11 years in the Adolescent Brain Cognitive Development (ABCD) dataset [87]. Our envirotyping data included nine youth-report social environment quality measures: Community Risk, Family Conflict, School Involvement, Neighborhood Safety, Caregiver Acceptance, School Disengagement, Parental Monitoring, School Environment, and Social Resilience. We used these specific measures because they represented all of the youth-report measures of social environment quality with at least an 80% response rate at the baseline time point. While measures such as Youth Prosocial Behavior, Discrimination, the Family Support subscale of the Mexican American Cultural Values measure, Involvement with Prosocial Peers, Involvement with Rule-breaking/Delinquent Peers, and Peer Network Health, among others, captured social environment quality, most were not recorded at baseline. We note that while objective measures like socioeconomic status have been widely used to examine brain-environment associations (e.g. 26, 65), recent work has suggested that subjective indicators of social environment quality may have greater bearing on mental health and wellbeing [17, 21]. Moreover, it is of theoretical interest to investigate youth’s own perceptions of their environments. All measures were z-scored to be on the same scale.

In our longitudinal analysis, we used a subset of the same social environment measures, which represented all of the youth-report subjective measures of social environment quality with at least an 80% response rate across three time points. That is, measures without sufficient response rate across all time points were dropped. Time points included baseline, 2-year follow-up, and 3-year follow-up. We opted to exclude the 1-year follow-up because all but one of our original measures were not recorded. All measures were z-scored independently at each time point to be on the same scale.

### K. Acquisition and pre-processing of neuroimaging data

Details of MRI acquisition for ABCD data have been described elsewhere [14]. Scans were typically completed on the same day as the social environment surveys, but could also be completed at a second testing session. After completing motion compliance training in a simulated scanning environment, subjects first underwent a structural T1-weighted scan. Then, subjects completed two five-minute resting-state scans, during which they were instructed to lay with their eyes open while a crosshair was on the screen. After the two resting-state scans, subjects completed two other structural scans as part of the larger ABCD protocol, followed by one or two more resting-state scans, depending on the protocol at the specific study site. All scans were collected on one of three 3T scanner platforms with an adult-size head coil.

The functional MRI data we used were minimally pre-processed according to the HCP minimal pre-processing pipeline described in [32]. Briefly, this pipeline includes distortion correction and alignment, denoising with Advanced Normalization Tools (ANTS89), FreeSurfer90 segmentation, surface registration, and volume registration using FSL FLIRT rigid-body transformation. Additional processing was done according to the DCAN BOLD Processing (DBP; 27) pipeline which included the following steps: (1) DBP standard pre-processing, (2) removing respiratory signal from motion realignment data by filtering out frequencies between 18.582 - 25.726 breaths per minute, (3) applying DPB motion censoring (frames exceeding an FD threshold of 0.2mm or failing to pass outlier detection at *±*0.3 standard deviations were discarded), and (4) DBP generation of parcellated time series into the Gordon 333 cortical and 19 subcortical atlas [35]. After quality control of the imaging data, we were left with 2776 adolescents (for a comparison of social environment scores from those with versus without quality imaging see Supplementary Figure S1).

### L. Construction of functional connectivity brain networks

Using the parcellated time series for 352 cortical and subcortical regions, we computed for each subject a functional connectivity brain network. To do this, we computed the Pearson correlation between the time series of all pairs of nodes for each subject. This procedure yielded a 352 *×* 352 matrix whose *ij*th entry was the Pearson correlation coefficient between nodes *i* and *j*, or the edge weight between nodes *i* and *j* in a functional connectivity (FC) brain network.

### M. Identifying envirotypes using hierarchical clustering

To identify envirotypes, we used a hierarchical clustering algorithm applied to the social environment data, which allowed us to tune the resolution of the clusters. To achieve the baseline clustering, we first computed the pairwise correlation matrix between all pairs of participants on the nine baseline measures of social environment quality, yielding a similarity matrix defined by social environment quality. We applied an arctangent transformation to this similarity matrix so all values were greater than 0.

To recursively partition this similarity matrix (and thereby participants) into clusters, we used a parameter-free hierarchical modularity maximization algorithm [7]. In general, the modularity, *Q*, of a partition can be expressed as the sum of contributions made by each community, *c ∈* 1, …, *K*, such that:

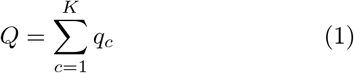

Where

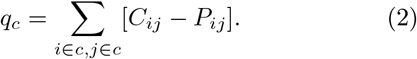

In this expression, *i* and *j* correspond to distinct elements in the matrix (i.e., distinct participants). The values of *C*_*ij*_ and *P*_*ij*_ correspond to the observed and expected correlation between those pairs of subjects, respectively. For a given correlation matrix, we uniformly set the expected weight of connections equal to the mean correlation value. That is 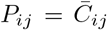 for all *i, j* pairs, where 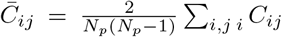 and *N*_*p*_ is participants. This pro the total number of cedure yields clusters of participants whose similarity to each other maximally exceeds what would be expected by chance. Because the algorithm operates recursively, there is no one “true” scale on which the resulting clusters are defined. This formalization allows us to zoom in to different resolutions of clusters to examine increasingly homogeneous envirotypes.

Our clustering algorithm is guaranteed to identify clusters, but it does not guarantee that the resulting clusters will be different from each other in a meaningful way. To characterize how exactly clusters differed from each other, we focus on Baseline Clusters 1 and 2 from hierarchical level 2, which together account for 89.3% of all participants. We performed a *t*-test to determine the extent to which these two clusters differed from each other across the nine measures. Additionally, we considered whether socioeconomic status (determined using a composite measure), pubertal status, age, or data collection site could account for cluster composition.

### N. Validating clusters

To validate that our identified clusters were meaningfully representative of their envirotypes—and not sensitive to the particular individuals included in our clustering algorithm—, we performed a subsampling procedure to assess the stability of cluster composition. To do this, we randomly sampled 10%, 20%, 30%, 40%, 50%, 60%, 70%, 80%, and 90% of the data. We clustered each subsample nonrecursively, so that we could compare the resulting cluster assignments to the hierarchical level 2 clusters. For each subsample, we computed for each individual *i* the percent of overlap of between people assigned to the same cluster as *i* in the subsample and original baseline clusters. For all subsample sizes, the mean percent overlap was *>* 0.875% (see Figure 1f). Additionally, we computed the percent overlap between the baseline and longitudinal clusters (described below).

After computing subsample clusters, we assessed how well the subsample cluster centroids aligned with the centroids of our identified baseline clusters. To do this, we used the Hungarian algorithm for the linear assignment problem. This procedure returns optimal pairs of clusters between the subsampled clusters and identified clusters. For the focal clusters (i.e., Baseline Clusters 1 and 2), we computed the correlation across scores between pairs (see Supplementary Figure S4).

### O. Examining differences in brain network organization between envirotypes

We subsequently compared these two focal clusters on the basis of their whole-brain functional connectivity (FC; N = 333 cortical and 19 subcortical regions of interest using the Gordon parcellation 34). First, we performed mass *t*-tests on the edge weights (i.e., the correlation coefficients of the time series of the nodes), correcting for multiple comparisons. Then, we evaluated statistical differences between clusters at the level of brain systems accounting for the spatial embedding of the brain using a ‘spin’ test. Briefly, this involved computing the mean difference in FC between every pair of brain systems and comparing this value against a null distribution generated using a space-preserving null model. FC differences were statistically significant if the observed difference exceeded that of the null distribution (false discovery rate fixed at *q* = 0.05; adjusted critical value of *p*_*crit*_ = 0.02).

### P. Quantifying social envirotype dynamics and associated outcomes

Since adolescence is a transitional period involving rapid change, we leveraged the longitudinality of the ABCD dataset to characterize both how envirotypes change over time and how these patterns of change are associated with diverging outcomes. We began by implementing the same procedure as in our first analyses. Namely, we computed the pairwise correlation of participants (*N* = 1529 with complete data) across 18 measures of social environment quality (i.e., six measures across three time points). Then we applied our hierarchical clustering algorithm, which yielded longitudinal clusters of participants at different levels of resolution whose similarity to each other maximally exceeds what would be expected by chance. As noted above, the recursivity of the algorithm entails there is no one “true” scale of the resulting clusters, which allows us to zoom in and out of different resolutions.

Finally, we examined how the dynamic envirotypes score on a suite of 28 outcome measures capturing cognitive, clinical, behavioral, and life experience variables. We selected measures across a wide range of categories that had at least an 80% response rate by our sample for the last time point. To determine whether the envirotypes scored differently from one another, we first performed a one-way ANOVA. For outcome measures for which there was a significant difference, we performed post-hoc Tukey’s tests. Then, to determine whether the longitudinal clusters were enriched with particular scores, we used permutation testing, which involved 1000 iterations of shuffling cluster labels, calculating mean cluster score for each outcome measure, and computing the difference between each permuted score and the observed score. We obtained *p*-values by dividing the number of instances in which the permuted difference was more extreme than the observed by the number of total permutations.

## SUPPLEMENTARY MATERIALS

**Table S1.** Summary of Measures for Clustering. T1 means and standard deviations are calculated using pre-z-scored data from all subjects included in baseline clustering (*N* = 2776), while T2 and T3 means and standard deviations are calculated using pre-z-scored data from only those included in longitudinal clustering (*N* = 1529).

| Measure | T1 Mean±SD (Min - Max) | T2 Mean±SD (Min - Max) | T3 Mean±SD (Min - Max) |
| --- | --- | --- | --- |
| Community Risk | 0.649 ± 0.838(0 – 4) | NA | NA |
| Caregiver Acceptance | 2.779 ± 0.281(1.2 – 3) | NA | NA |
| Neighborhood Safety | 4 ± 1.016(1 – 5) | NA | NA |
| Parental Monitoring | 4.423 ± 0.479(2.2 – 5) | 4.513 ± 0.447(1.8 – 5) | 4.418 ± 0.478(2 – 5) |
| Family Conflict | 2.014 ± 1.894(0 – 9) | 1.825 ± 1.818(0 – 9) | 1.999 ± 1.962(0 – 9) |
| School Environment | 19.84 ± 2.588(8 – 24) | 19.73 ± 2.762(9 – 24) | 19.6 ± 2.701(8 – 24) |
| School Involvement | 13.06 ± 2.187(4 – 16) | 12.76 ± 2.36(4 – 16) | 12.66 ± 2.212(4 – 16) |
| School Disengagement | 3.793 ± 1.381(2 – 8) | 3.902 ± 1.358(2 – 8) | 4.094 ± 1.299(2 – 8) |
| Social Resilience | 6.309 ± 7.585(0 – 143) | 6.788 ± 7.71(0 – 105) | 6.403 ± 6.911(0 – 150) |

**Table S2.** Summary of Measures for Composite Socioeconomic Status.

| Measure | Mean±SD (Min - Max) |
| --- | --- |
| Parent Employment | 2.268 ± 2.276(1 – 11) |
| Partner Employment | 1.495 ± 1.579(1 – 11) |
| Parent Education | 16.89 ± 2.54(1 – 21) |
| Partner Education | 16.66 ± 2.814(1 – 21) |
| Parent Income | 5.083 ± 2.693(1 – 10) |
| Partner Income | 6.902 ± 2.337(1 – 10) |
| Combined Income | 7.553 ± 2.199(1 – 10) |
| Number of People in Home | 4.679 ± 1.545(0 – 16) |
| Afford Food | 0.069 ± 0.254(0 – 1) |
| Afford Rent | 0.096 ± 0.294(0 – 1) |
| Afford Phone | 0.051 ± 0.219(0 – 1) |
| Afford Utilities | 0.049 ± 0.215(0 – 1) |
| Afford Doctor | 0.052 ± 0.221(0 – 1) |
| Afford Dentist | 0.099 ± 0.299(0 – 1) |
| Been Evicted | 0.01 ± 0.099(0 – 1) |

**Table S3.** Summary of Outcome Measures.

| Measure | Mean±SD (Min - Max) |
| --- | --- |
| CBCL Attention | 53.37 ± 5.242(50 – 95) |
| CBCL Aggressive | 52.28 ± 4.691(50 – 86) |
| CBCL Rulebreak | 51.89 ± 3.743(50 – 78) |
| CBCL Social Problems | 52.41 ± 4.601(50 – 88) |
| CBCL Thought Problems | 53.54 ± 5.629(50 – 80) |
| CBCL Anxiety and Depression | 53.66 ± 6.048(50 – 89) |
| CBCL Withdrawal, Depression | 53.66 ± 5.787(50 – 90) |
| CBCL Depression | 53.96 ± 6.104(50 – 89) |
| CBCL Somatic Problems | 54.47 ± 6.101(50 – 93) |
| CBCL Oppositional Defiance Disorder | 52.42 ± 4.765(50 – 80) |
| CBCL Anxiety Disorder | 53.63 ± 6.219(50 – 94) |
| CBCL Somatic | 54.38 ± 5.903(50 – 96) |
| CBCL Conduct Disorder | 52.22 ± 4.408(50 – 77) |
| CBCL Stress | 53.12 ± 5.626(50 – 85) |
| Prosocial Behavior | 1.682 ± 0.433(0 – 2) |
| Positive Affect | 20.23 ± 4.059(9 – 27) |
| Impulsivity | 2.631 ± 6.127(0 – 58) |
| Bad Experiences | 2.177 ± 2.121(0 – 13) |
| Good Experiences | 1.608 ± 1.407(0 – 8) |
| Affected by Bad Experiences | 4.396 ± 4.883(0 – 6) |
| Affected by Good Experiences | 2.907 ± 3.173(0 – 5) |
| Peer Experiences: Relational Victim | 4.695 ± 1.846(3 – 13) |
| Peer Experiences: Relational Aggression | 3.908 ± 1.298(3 – 12) |
| Peer Experiences: Overt Victim | 3.535 ± 1.189(3 – 15) |
| Peer Experiences: Overt Aggression | 3.236 ± 0.753(3 – 11) |
| Peer Experiences: Reputation Victim | 3.95 ± 1.759(3 – 15) |
| Peer Experiences: Reputation Aggression | 3.167 ± 0.622(3 – 13) |
| Problem Solving | 3.817 ± 0.748(1 – 5) |

**Figure S1.**
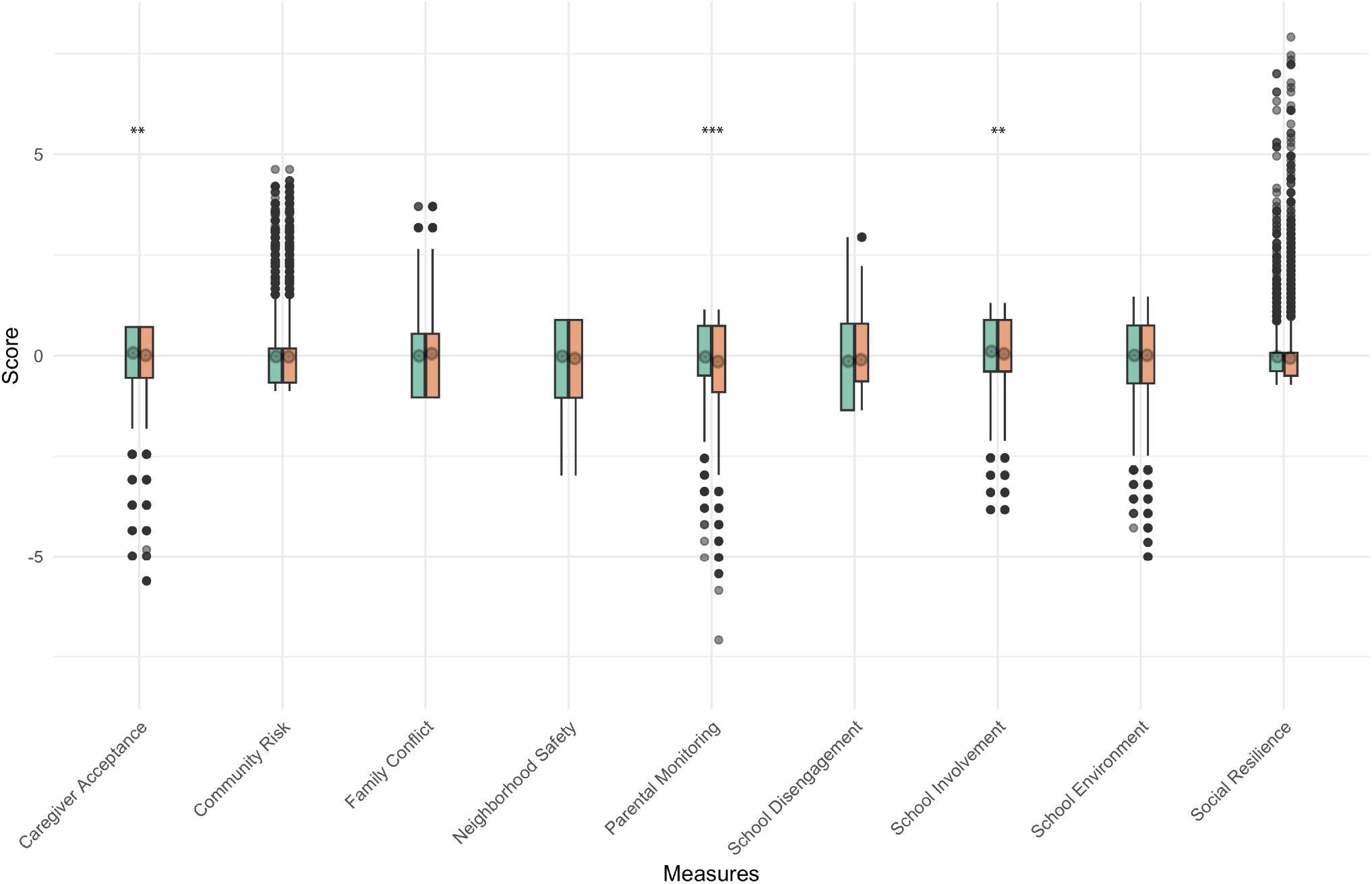
Comparing social environment scores of those with versus without quality imaging data. To quantify how representative our clusters are of the entire dataset, for each measure used to cluster people into envirotypes, we compared the scores of those with complete quality data who were included in the clustering (blue boxes on left) to all other subjects in the data (pink boxes on right). While there were significant differences between the groups on three of the nine measures (Caregiver Acceptance, Parental Monitoring, and School Involvement; *p*s *<* 0.01), the effect sizes were not especially large and the remaining measures showed no difference. Asterisks indicate significance level, such that ** means *p <* 0.01 and *** means *p <* 0.001.

**Figure S2.**
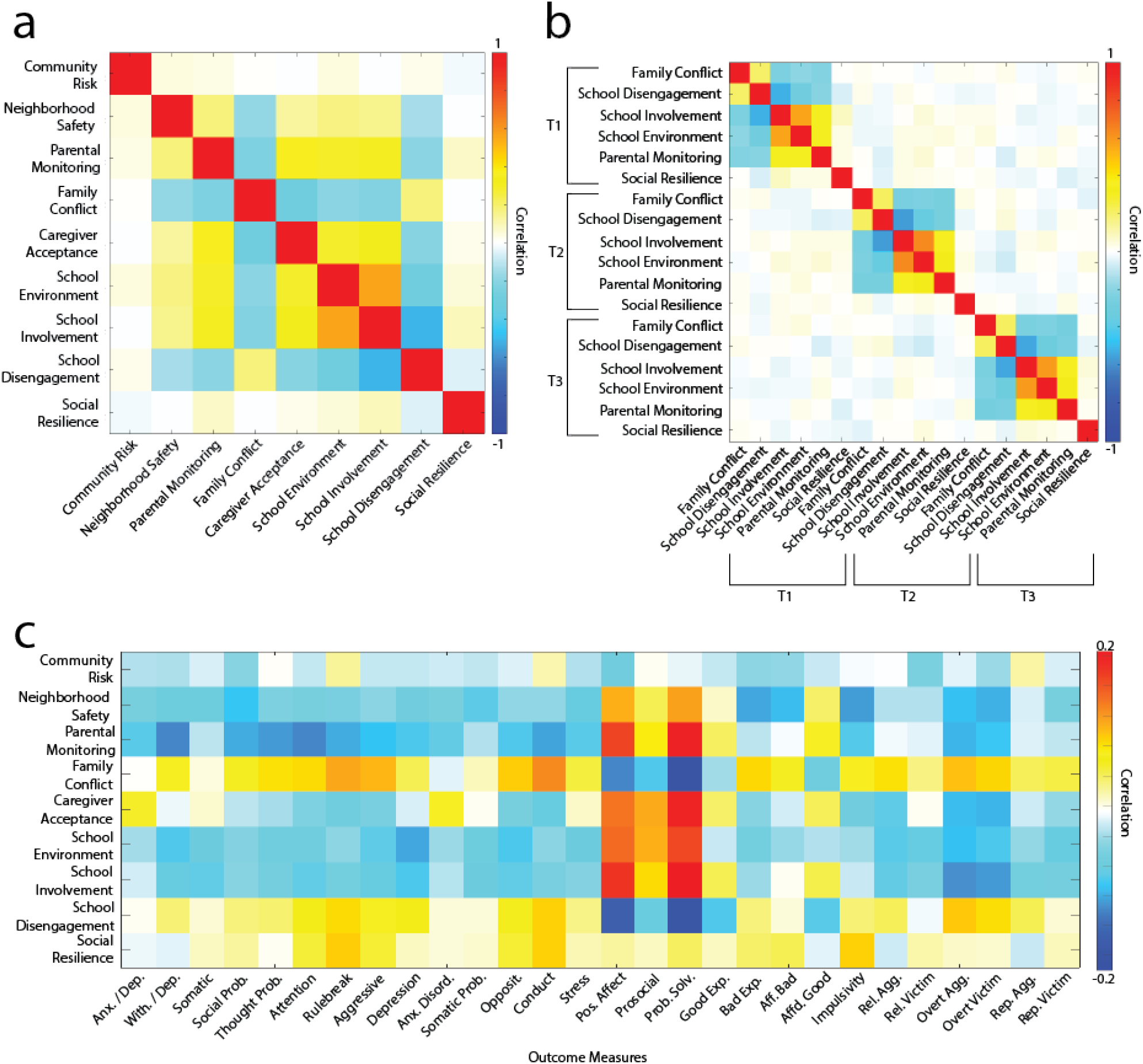
Correlations between measures used for clustering. Panel (a) shows the correlations between the measures used for baseline envirotype clustering. Panel (b) shows the correlations between measures used for longitudinal envirotype clustering. Panel (c) shows the correlation between clustering measures at baseline and the outcome measures.

**Figure S3.**
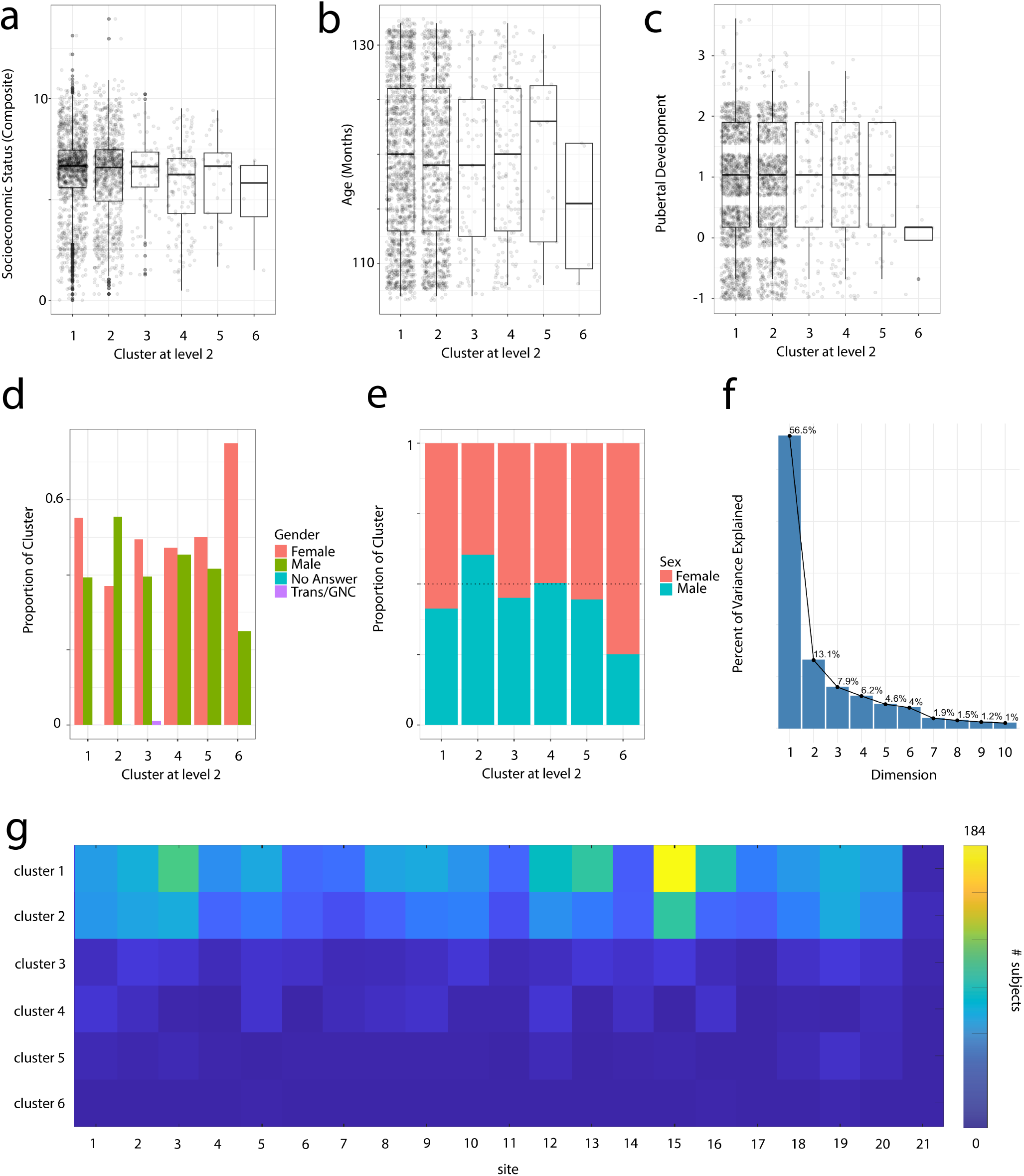
Distributions of demographic variables across clusters. Panel (a) shows the distribution of a composite measure of socioeconomic status across clusters. The only significant difference, identified using an ANOVA and post-hoc Tukey’s test, was between Cluster 1 and Cluster 4 (*p* = 0.007). Panel (b) shows the distribution of age in months across clusters. There were no differences in age between clusters according to an ANOVA. Panel (c) shows the pubertal development scores across clusters. There were no differences in pubertal development between clusters according to an ANOVA. Panels (d) and (e) provide the distributions of gender and sex, respectively, across clusters. Panel (f) gives the percent of variance explained by the first 10 dimensions identified through PCA applied to the 15 individual measures of socioeconomic status (see Table S2 for descriptive statistics of these measures). Coefficients from the first principal component were used to compute the composite socioeconomic status score as shown in panel (a). Panel (g) shows the number of subjects from each data acquisition site in each cluster.

**Figure S4.**
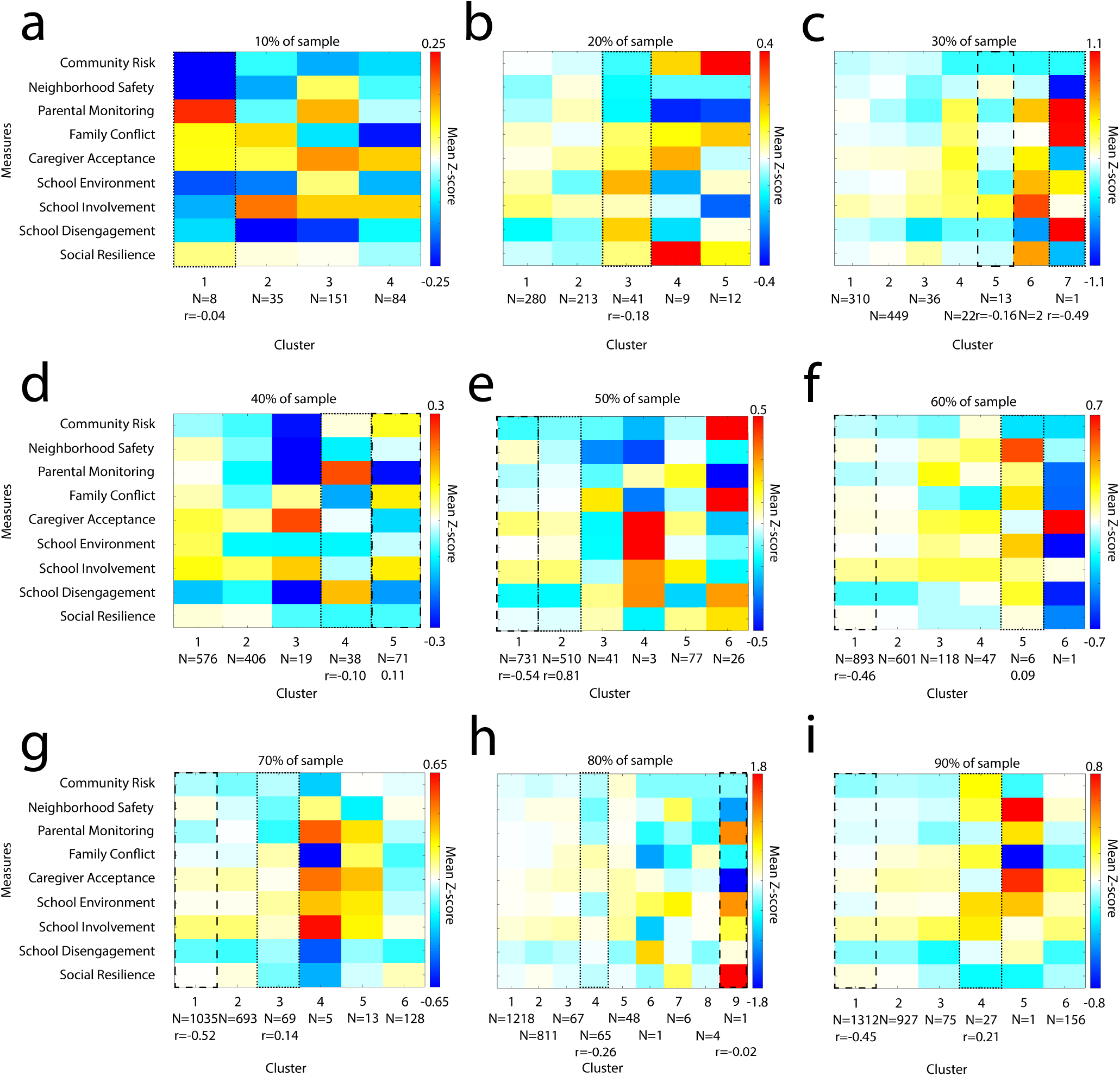
Subsampling centroids. We show the centroids for all clusters for each subsample. We matched all subsampled clusters to observed clusters using the Hungarian algorithm for the linear assignment problem and note both the matches for focal clusters 1 (dotted outline) and 2 (dashed outline) described in the main text and the correlations between matched clusters across measures. We note that cluster 3 did not have a match when we subsampled with 10% or 20% of data.

**Figure S5.**
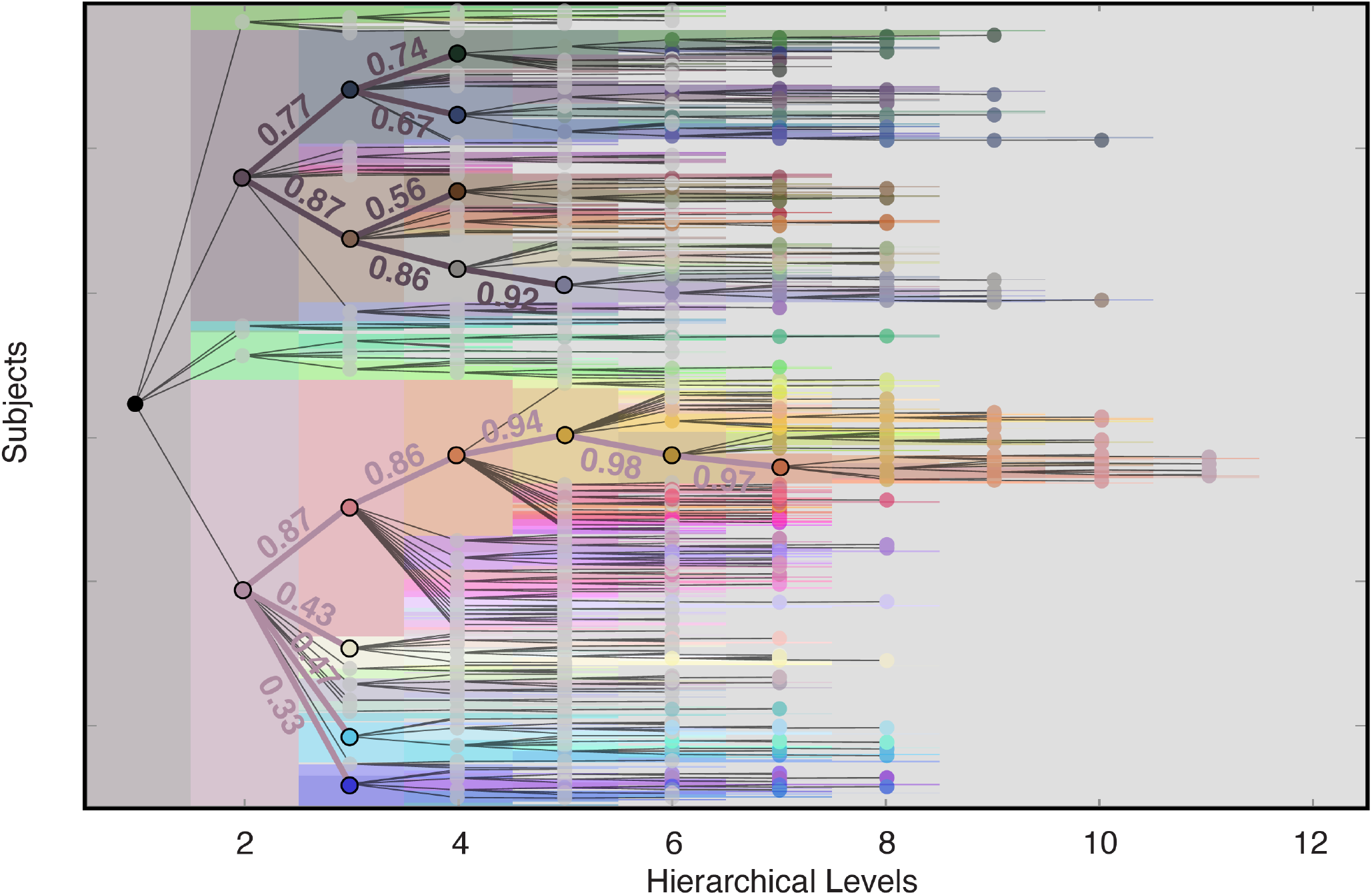
Baseline cluster refinement. We compare more fine grain clusters to their coarser parents across hierarchical levels. We compute the correlation between centroids (mean scores across all cluster members for each measure) of child clusters to their level 2 parent. We opt to compare to level 2 parents, since we focused on this resolution in the main text. Additionally, we only include child clusters with at least 100 members. A dendrogram of cluster inheritance is overlaid atop the figure of hierarchical levels from Figure 1c for clarity. Nodes in the dendrogram identify clusters, and edges indicate a parent-child relationship. Nodes of clusters we compared to coarser parent centroids are outlined in black. Edges and the text of the parent-child correlations are colored to match the level two cluster to which they belong.

**Figure S6.**
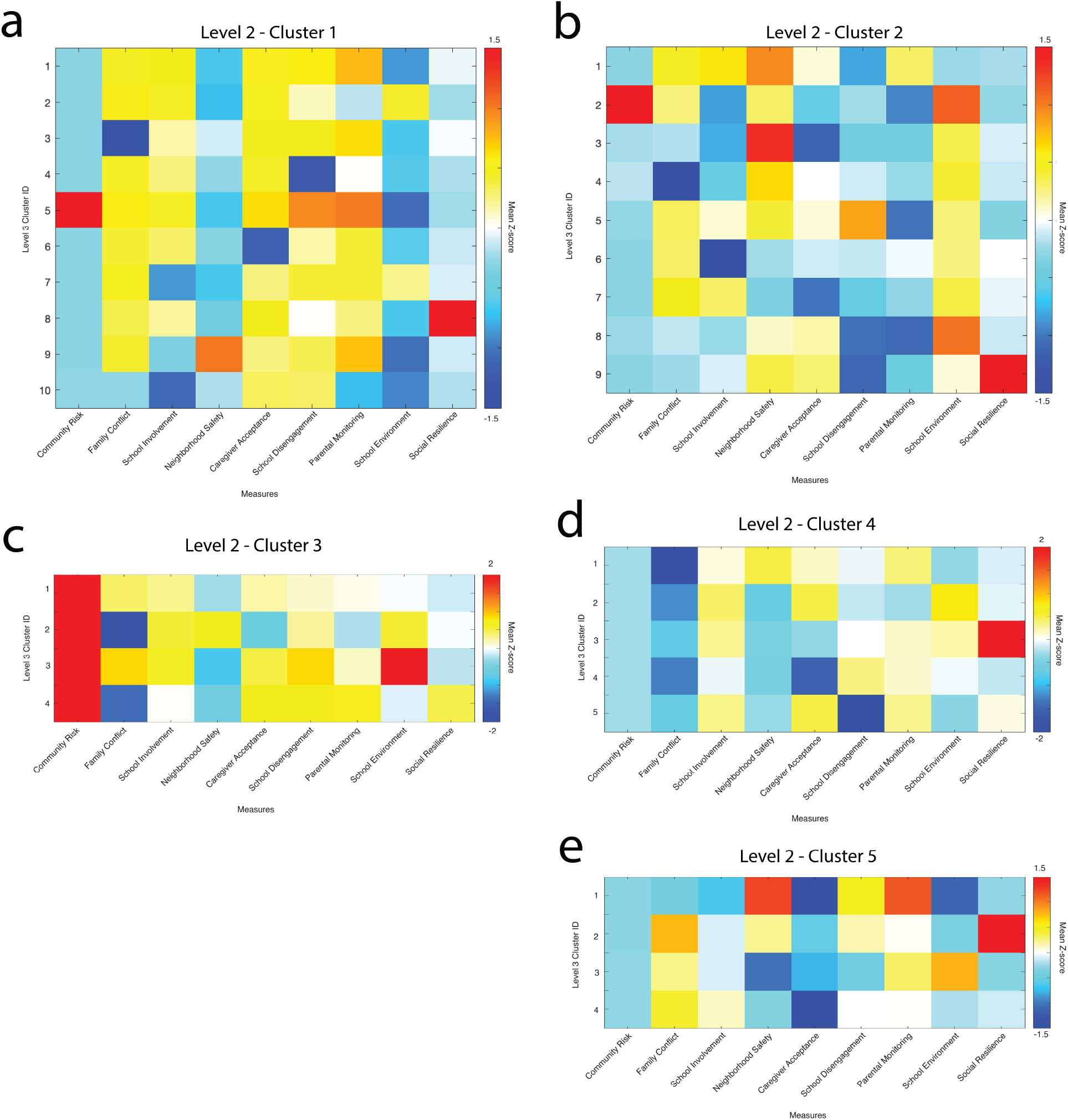
Envirotypes for Level 3 Clusters. Here we show how the level 2 clusters differentiate at a finer resolution. Cluster 1 on level 2 separated into 10 clusters on level 3, with *N*^1^ = 890, *N*^2^ = 106, *N*^3^ = 140, *N*^4^ = 33, *N*^5^ = 31, *N*^6^ = 34, *N*^7^ = 77, *N*^8^ = 31, *N*^9^ = 110, and *N*^10^ = 5 subjects and with 18 singletons. Cluster 2 on level 2 separated into 4 clusters on level3, with *N*^1^ = 11, *N*^2^ = 6, *N*^3^ = 3, and *N*^4^ = 8 and with 13 singletons. Cluster 3 on level 2 separated into 9 clusters on level 3, with *N*^1^ = 9, *N*^2^ = 16, *N*^3^ = 444, *N*^4^ = 54, *N*^5^ = 7, *N*^6^ = 64, *N*^7^ = 17, *N*^8^ = 391, and *N*^9^ = 6 and with 2 singletons. Cluster 4 on level 2 separated into 5 clusters on level 3, with *N*^1^ = 69, *N*^2^ = 52, *N*^3^ = 24, *N*^4^ = 13, and *N*^5^ = 3 and with 4 singletons. Cluster 5 on level 2 separated into 4 clusters on level 3, with *N*^1^ = 3, *N*^2^ = 8, *N*^3^ = 3, and *N*^4^ = 21 and with 1 singleton.

**Figure S7.**
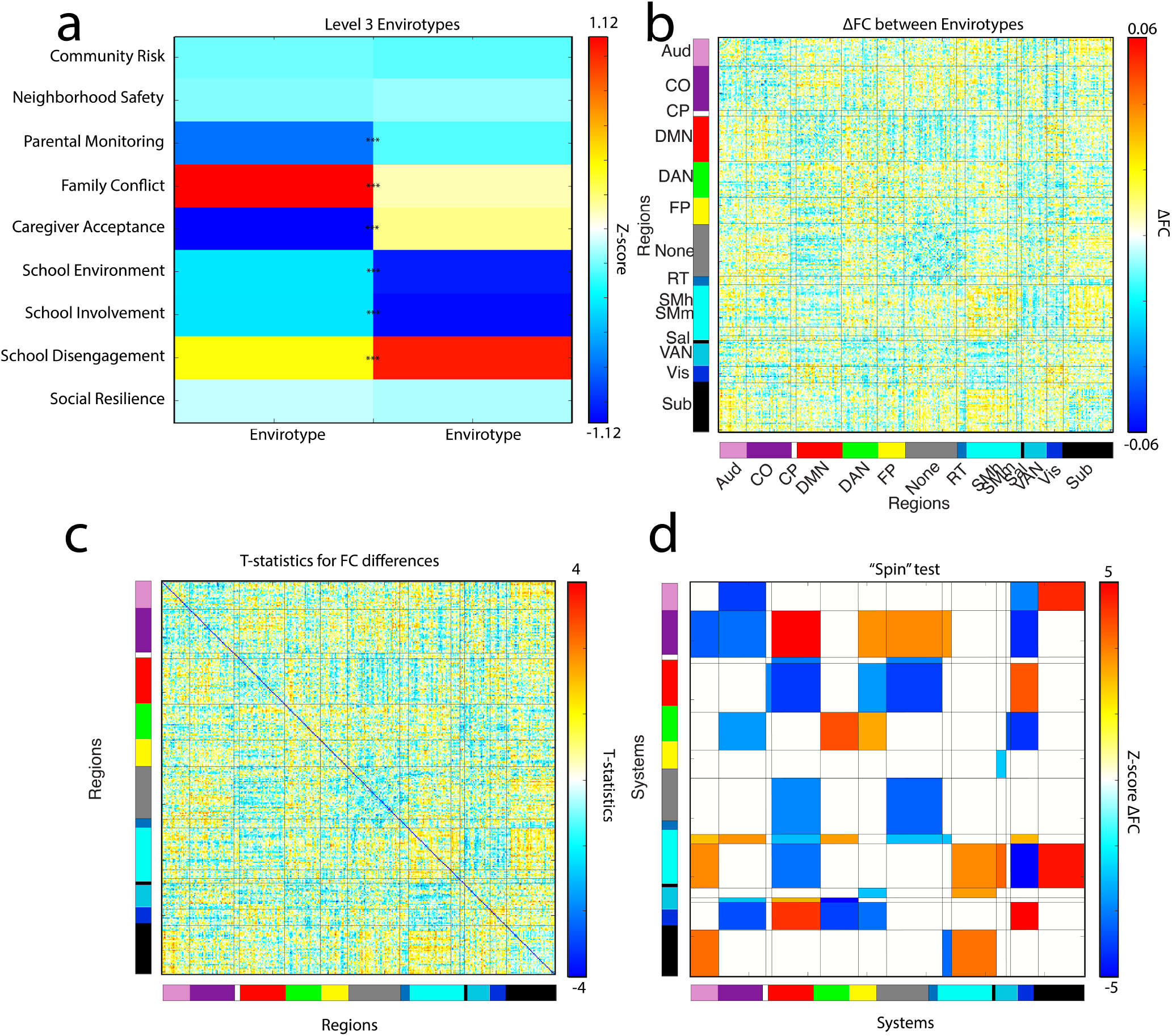
Comparing Level 3 Clusters. Here we compare the envirotypes and brain network organization of the two large clusters that appear in level 3 (in red and green in Figure 1c) as the lower quality envirotype in level 2 (in brown in Figure 1c) fractures with increased resolution. These envirotypes are also shown in Figure S6 as part of cluster 2. Panel (a) shows the envirotypes for these clusters, which different significantly on 6 out of 9 measures (Parental Monitoring, Family Conflict, Caregiver Acceptance, School Environment, School Involvement, and School Disengagement; all *t* s *>* 7.6, all *p*s *<* 0.0001). Panel (b) shows the difference in each cluster’s mean functional connectivity, the |*t*|-statistics for which are in panel (c). Additionally, we performed space-preserving “spin” tests, shown in panel (d).

**Figure S8.**
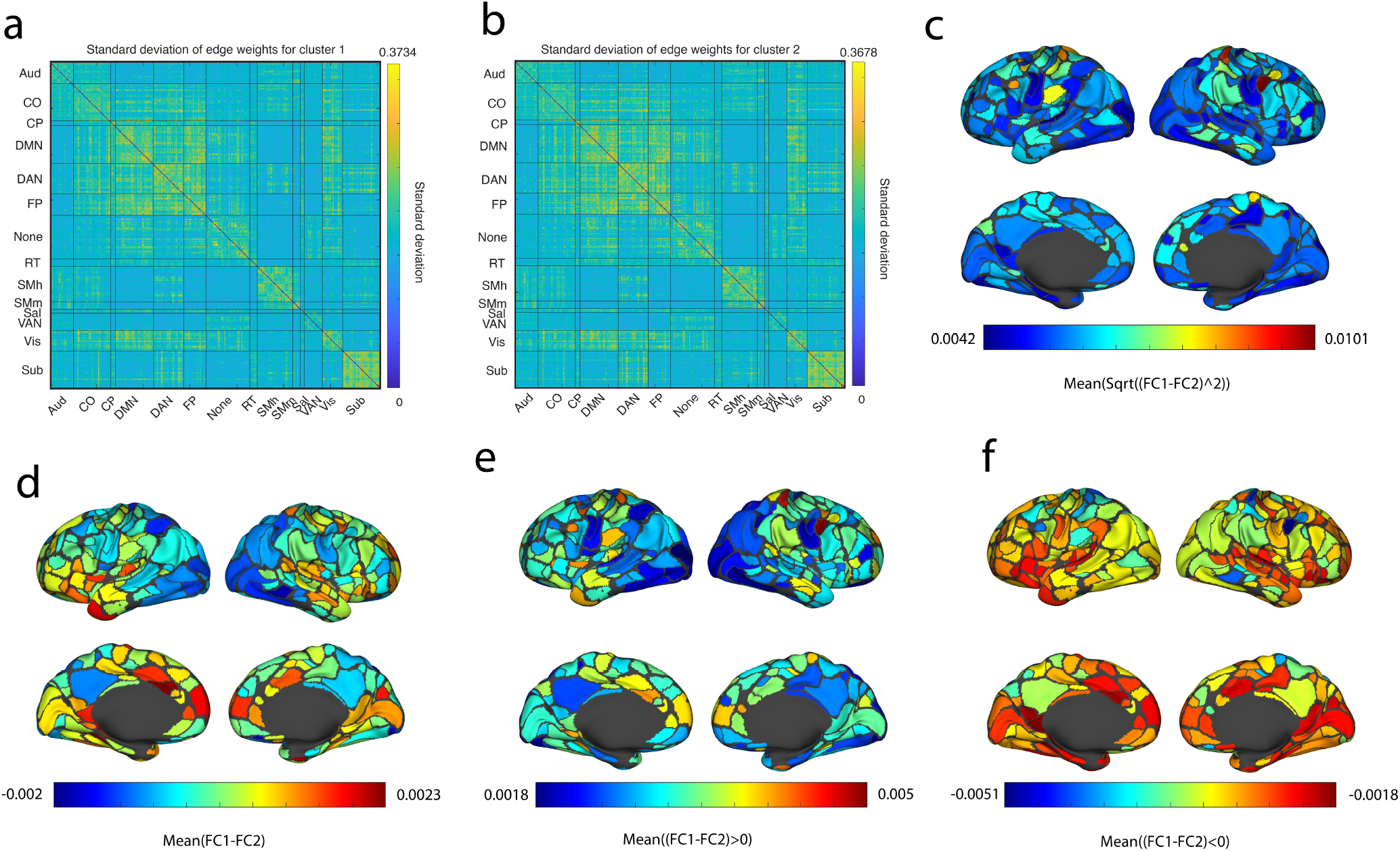
Differences in FC between Clusters 1 and 2. Panels (a) and (b) show the standard deviations of edge weights for members of clusters 1 and 2, respectively. Panel (c) shows mean square root of the squared difference in FC between clusters, which makes clear the magnitude of the differences between clusters. Panel (d) shows the mean difference in FC between clusters, which emphasizes the sign of the differences between clusters. Panels (e) and (f) separate these differences by sign, showing the mean of only the positive and negative differences, respectively. When the difference is positive, cluster 1–which is characterized by a “higher quality” envirotype has greater edge weights; when the difference is negative, cluster 2 has greater edge weights.

**Figure S9.**
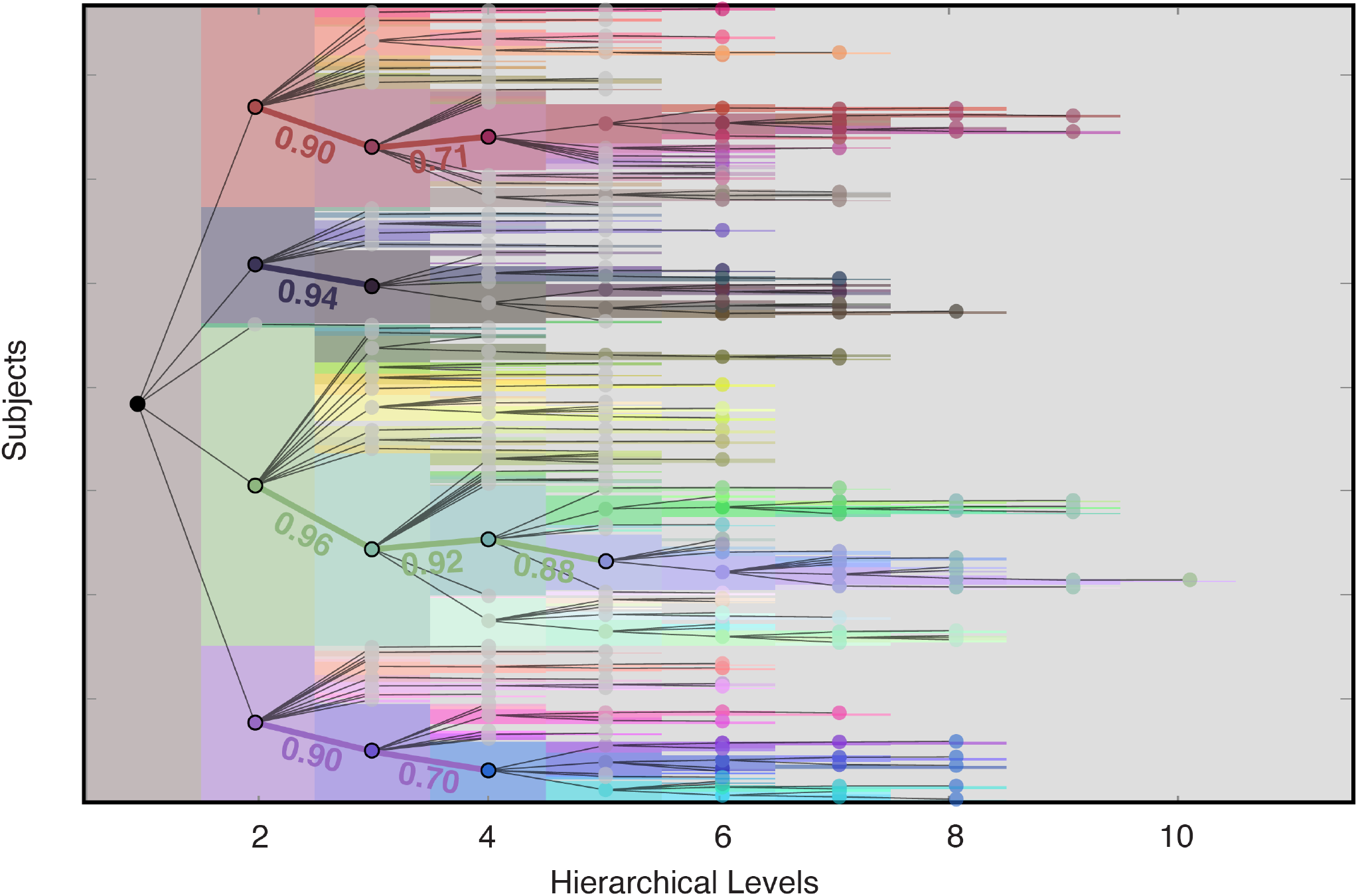
Refinement of longitudinal clusters. We examine how the longitudinal clusters refine over hierarchical levels in the same way as the baseline clusters. That is, we compute the correlation between centroids (mean scores across all cluster members for each measure) of child clusters to their level 2 parent. Again, we compare to level 2 parents since we focused on this resolution in the main text and we only include clusters with at least 100 members. A dendrogram of cluster inheritance is overlaid atop the figure of longitudinal hierarchical clusters from Figure 33 for clarity. Nodes in the dendrogram identify clusters, and edges indicate a parent-child relationship between clusters. Nodes of clusters we compared to coarser parent centroids are outlined in black. Edges and the text of the parent-child correlations are colored to match the level 2 cluster to which they belong.

**Figure S10.**
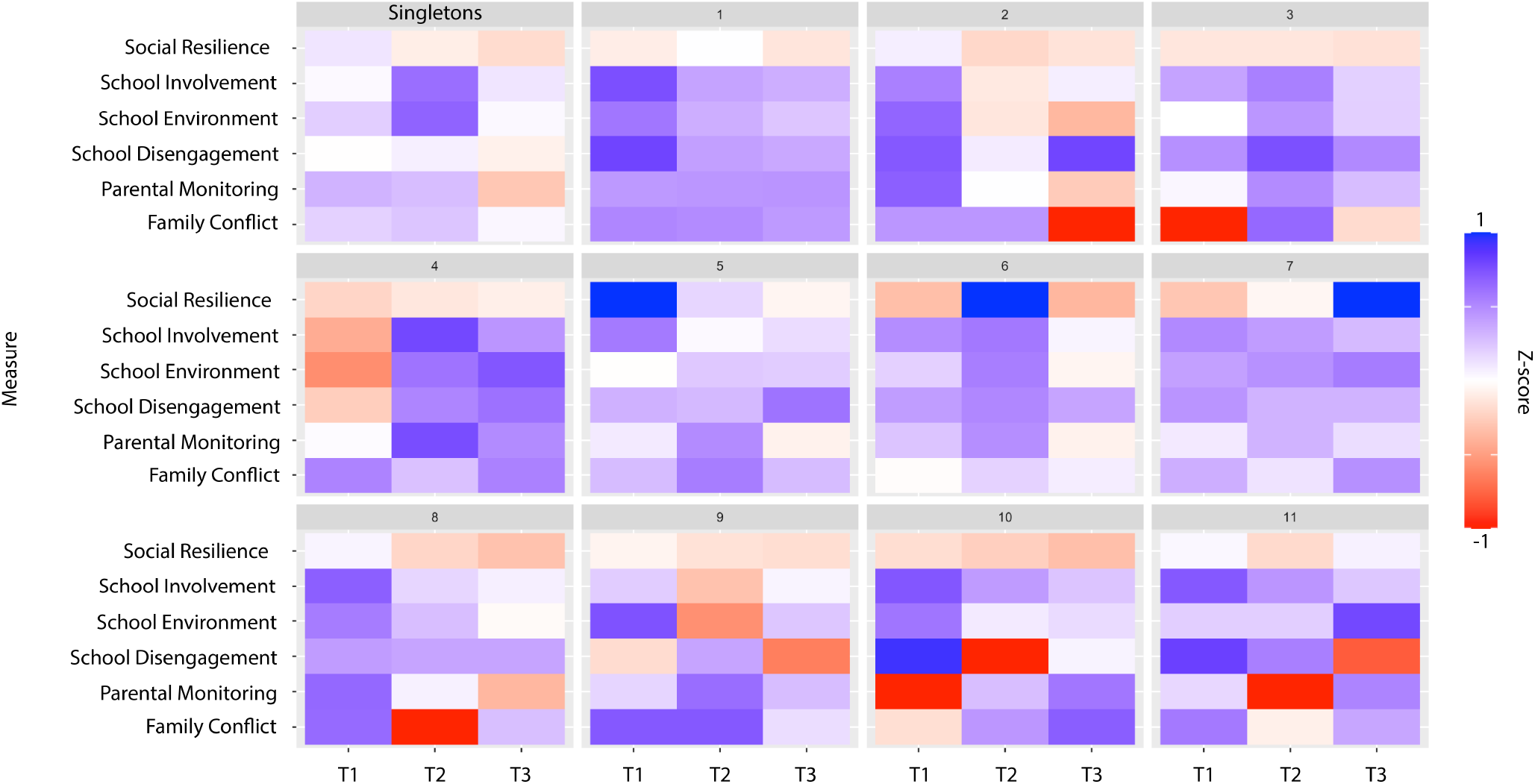
Cluster 1 finer resolution social envirotypes over time. Panels (a-f) show how finer resolution longitudinal envirotypes change over time for each of the six measures. All subjects shown here were assigned to cluster 1 on level 2, which was characterized by a stable, high quality social environment. Cluster 1 differentiates into 11 clusters plus singletons, with cluster sizes *N*_1_ = 370, *N*_2_ = 9, *N*_3_ = 49, *N*_4_ = 51, *N*_5_ = 21, *N*_6_ = 23, *N*_7_ = 16, *N*_8_ = 20, *N*_9_ = 20, *N*_10_ = 17, and *N*_11_ = 8. There were 10 singletons.

**Figure S11.**
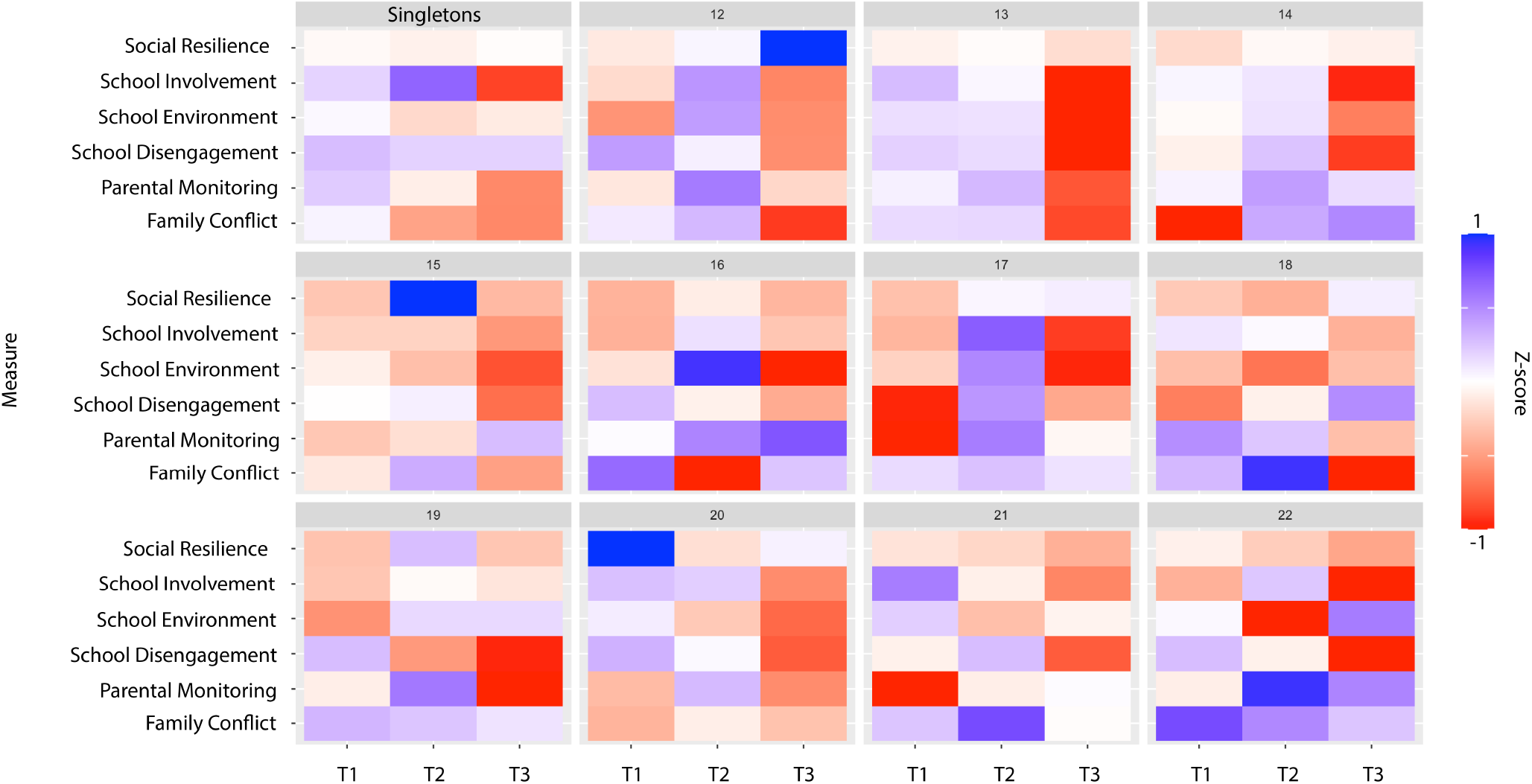
Cluster 2 finer resolution social envirotypes over time. Panels (a-f) show how finer resolution longitudinal envirotypes change over time for each of the six measures. All subjects shown here were assigned to cluster 2 on level 2, which was characterized by a worsening social environment by the last time point. Cluster 2 at level 2 differentiates into 11 clusters plus singletons at level 3, with cluster sizes *N*_12_ = 10, *N*_13_ = 191, *N*_14_ = 26, *N*_15_ = 10, *N*_16_ = 4, *N*_17_ = 19, *N*_18_ = 5, *N*_19_ = 8, *N*_20_ = 14, *N*_21_ = 4 and *N*_22_ = 2. There were 8 singletons.

**Figure S12.**
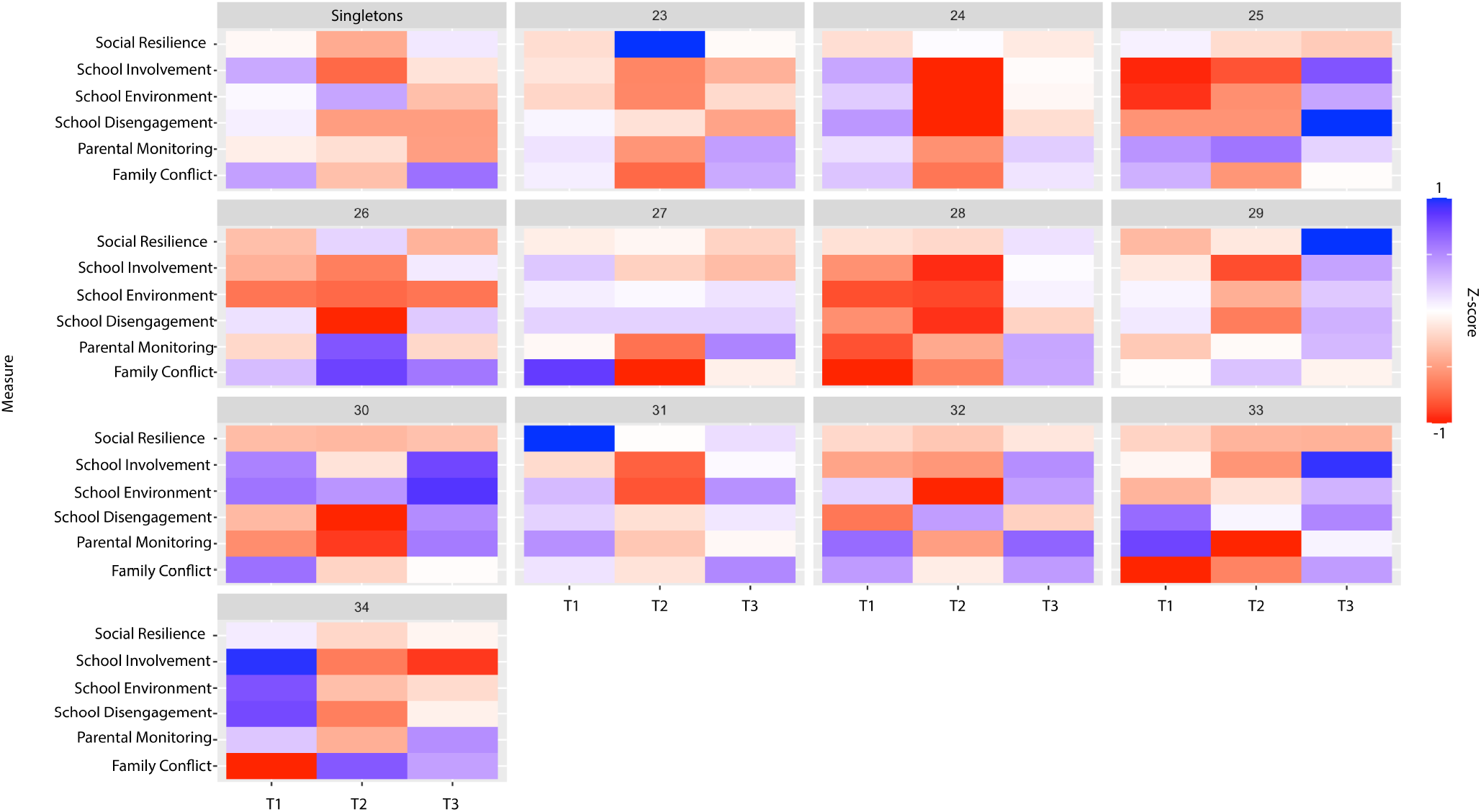
Cluster 3 finer resolution social envirotypes over time. Panels (a-f) show how finer resolution longitudinal envirotypes change over time for each of the six measures. All subjects shown here were assigned to cluster 3 on level 2, which was characterized by an unstable social environment with especially low quality in the second time point. Cluster 3 at level 2 differentiates into 11 clusters plus singletons at level 3, with cluster sizes *N*_23_ = 5, *N*_24_ = 229, *N*_25_ = 3, *N*_26_ = 7, *N*_27_ = 8, *N*_28_ = 56, *N*_29_ = 21, *N*_30_ = 5, *N*_31_ = 8, *N*_32_ = 13, *N*_33_ = 6 and *N*_34_ = 5. There were 5 singletons.

**Figure S13.**
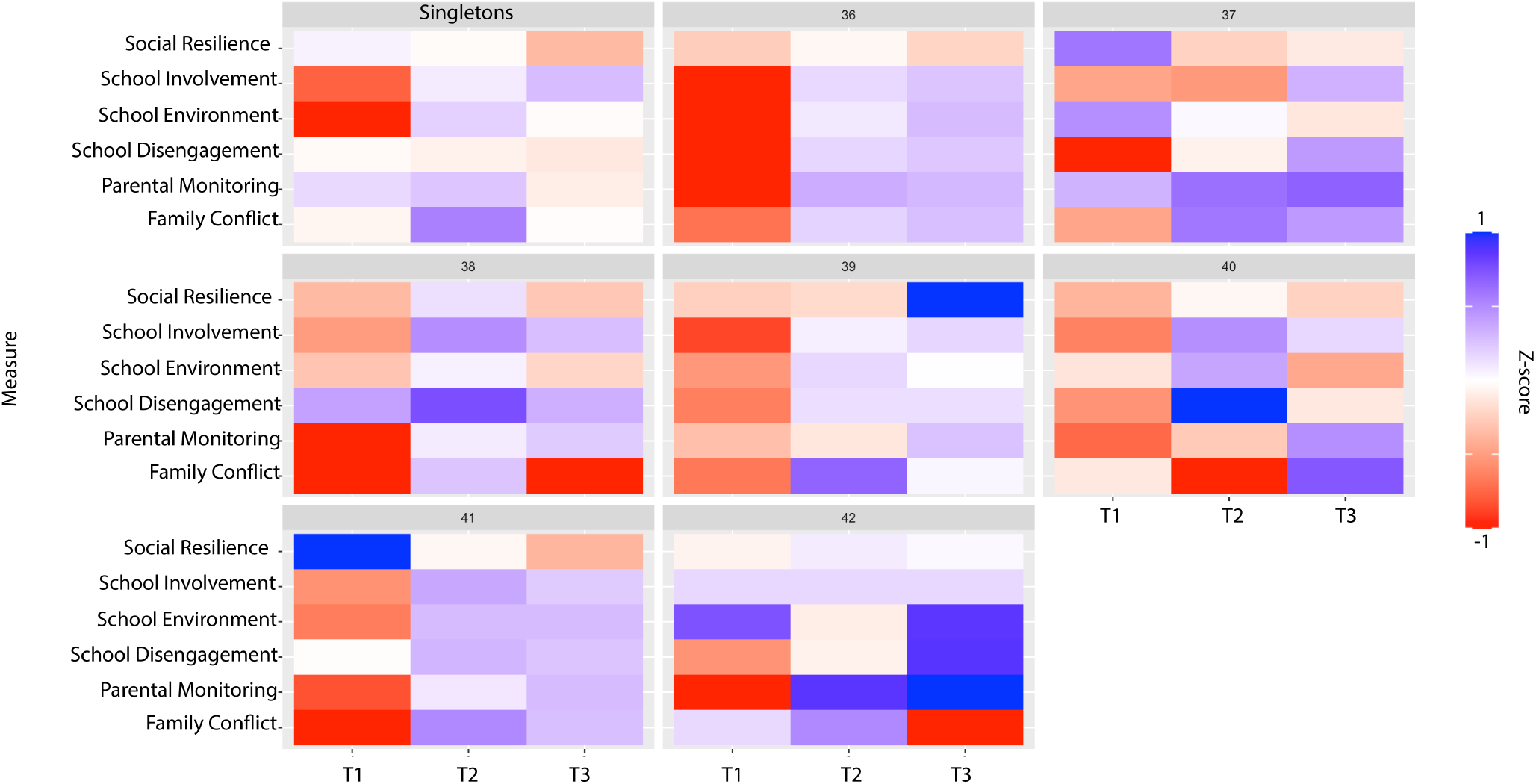
Cluster 4 finer resolution social envirotypes over time. Panels (a-f) show how finer resolution longitudinal envirotypes change over time for each of the six measures. All subjects shown here were assigned to cluster 4 on level 2, which was characterized by an improving social environment. Cluster 4 differentiates into 11 clusters plus singletons, with cluster sizes *N*_36_ = 140, *N*_37_ = 7, *N*_38_ = 12, *N*_39_ = 20, *N*_40_ = 15, *N*_41_ = 11, and *N*_42_ = 3. There were 15 singletons.

## References

[1] Astle, D. E., Bassett, D. S., and Viding, E. (2024). Understanding divergence: Placing developmental neuroscience in its dynamic context. Neuroscience & Biobehavioral Reviews, page 105539.

[2] Barch, D. M., Albaugh, M. D., Avenevoli, S., Chang, L., Clark, D. B., Glantz, M. D., Hudziak, J. J., Jernigan, T. L., Tapert, S. F., Yurgelun-Todd, D., et al. (2018). Demographic, physical and mental health assessments in the adolescent brain and cognitive development study: Rationale and description. Developmental cognitive neuroscience, 32:55–66.

[3] Baumrind, D. (1996). The discipline controversy revisited. Family relations, pages 405–414.

[4] Beckes, L. and Sbarra, D. A. (2022). Social baseline theory: State of the science and new directions. Current Opinion in Psychology, 43:36–41.

[5] Berkman, L. F. and Syme, S. L. (1979). Social networks, host resistance, and mortality: a nine-year follow-up study of alameda county residents. American journal of Epidemiology, 109(2):186–204.

[6] Betzel, R. F., Byrge, L., He, Y., Goñi, J., Zuo, X.-N., and Sporns, O. (2014). Changes in structural and functional connectivity among resting-state networks across the human lifespan. Neuroimage, 102:345–357.

[7] Betzel, R. F., Cutts, S. A., Tanner, J., Greenwell, S. A., Varley, T., Faskowitz, J., and Sporns, O. (2023). Hierarchical organization of spontaneous co-fluctuations in densely sampled individuals using fmri. Network Neuroscience, 7(3):926–949.

[8] Blanchett, R., Chen, Y., Aguate, F., Xia, K., Cornea, E., Burt, S. A., de Los Campos, G., Gao, W., Gilmore, J. H., and Knickmeyer, R. C. (2023). Genetic and environmental factors influencing neonatal resting-state functional connectivity. Cerebral Cortex, 33(8):4829– 4843.

[9] Boyce, W. T. and Ellis, B. J. (2005). Biological sensitivity to context: I. an evolutionary–developmental theory of the origins and functions of stress reactivity. Development and psychopathology, 17(2):271–301.

[10] Brakowski, J., Spinelli, S., Dörig, N., Bosch, O. G., Manoliu, A., Holtforth, M. G., and Seifritz, E. (2017). Resting state brain network function in major depression–depression symptomatology, antidepressant treatment effects, future research. Journal of psychiatric research, 92:147–159.

[11] Brody, G. H., Yu, T., Nusslock, R., Barton, A. W., Miller, G. E., Chen, E., Holmes, C., McCormick, M., and Sweet, L. H. (2019). The protective effects of supportive parenting on the relationship between adolescent poverty and resting-state functional brain connectivity during adulthood. Psychological science, 30(7):1040–1049.

[12] Bronfenbrenner, U. (1979). The ecology of human development: Experiments by nature and design. Harvard university press.

[13] Byrge, L., Sporns, O., and Smith, L. B. (2014). Developmental process emerges from extended brain– body–behavior networks. Trends in cognitive sciences, 18(8):395–403.

[14] Casey, B. J., Cannonier, T., Conley, M. I., Cohen, A. O., Barch, D. M., Heitzeg, M. M., Soules, M. E., Teslovich, T., Dellarco, D. V., Garavan, H., et al. (2018). The adolescent brain cognitive development (abcd) study: imaging acquisition across 21 sites. Developmental cognitive neuroscience, 32:43–54.

[15] Chan, M. Y., Na, J., Agres, P. F., Savalia, N. K., Park, D. C., and Wig, G. S. (2018). Socioeconomic status moderates age-related differences in the brain’s functional network organization and anatomy across the adult lifespan. Proceedings of the National Academy of Sciences, 115(22):E5144–E5153.

[16] Chen, E. and Miller, G. E. (2007). Stress and inflammation in exacerbations of asthma. Brain, behavior, and immunity, 21(8):993–999.

[17] Coan, J. A., Beckes, L., Gonzalez, M. Z., Maresh, E. L., Brown, C. L., and Hasselmo, K. (2017). Relationship status and perceived support in the social regulation of neural responses to threat. Social Cognitive and Affective Neuroscience, 12(10):1574–1583.

[18] Cohen, S. (1988). Psychosocial models of the role of social support in the etiology of physical disease. Health psychology, 7(3):269.

[19] Cohen, S. (2004). Social relationships and health. American psychologist, 59(8):676.

[20] Cohen, S. and Wills, T. A. (1985). Stress, social support, and the buffering hypothesis. Psychological bulletin, 98(2):310.

[21] Danese, A. and Widom, C. S. (2020). Objective and subjective experiences of child maltreatment and their relationships with psychopathology. Nature human behaviour, 4(8):811–818.

[22] De Jaegher, H., Di Paolo, E., and Gallagher, S. (2010). Can social interaction constitute social cognition? Trends in cognitive sciences, 14(10):441–447.

[23] Del Giudice, M., Ellis, B. J., and Shirtcliff, E. A. (2011). The adaptive calibration model of stress responsivity. Neuroscience & biobehavioral reviews, 35(7):1562–1592.

[24] Duque-Alarcón, X., Alcalá-Lozano, R., González-Olvera, J. J., Garza-Villarreal, E. A., and Pellicer, F. (2019). Effects of childhood maltreatment on social cognition and brain functional connectivity in borderline personality disorder patients. Frontiers in psychiatry, 10:156.

[25] Ellis, B. J., Boyce, W. T., Belsky, J., Bakermans-Kranenburg, M. J., and Van IJzendoorn, M. H. (2011). Differential susceptibility to the environment: An evolutionary–neurodevelopmental theory. Development and psychopathology, 23(1):7–28.

[26] Ellwood-Lowe, M. E., Whitfield-Gabrieli, S., and Bunge, S. A. (2021). Brain network coupling associated with cognitive performance varies as a function of a child’s environment in the abcd study. Nature Communications, 12(1):1–14.

[27] Feczko, E., Conan, G., Marek, S., Tervo-Clemmens, B., Cordova, M., Doyle, O., Earl, E., Perrone, A., Sturgeon, D., Klein, R., et al. (2021). Adolescent brain cognitive development (abcd) community mri collection and utilities. BioRxiv, pages 2021–07.

[28] Fortunato, S. and Barthelemy, M. (2007). Resolution limit in community detection. Proceedings of the national academy of sciences, 104(1):36–41.

[29] Foulkes, L. and Blakemore, S.-J. (2018). Studying individual differences in human adolescent brain development. Nature neuroscience, 21(3):315–323.

[30] Garavan, H., Bartsch, H., Conway, K., Decastro, A., Goldstein, R., Heeringa, S., Jernigan, T., Potter, A., Thompson, W., and Zahs, D. (2018). Recruiting the abcd sample: Design considerations and procedures. Developmental cognitive neuroscience, 32:16–22.

[31] Ghazanfar, A. A. and Gomez-Marin, A. (2024). The central role of the individual in the history of brains. Neuroscience & Biobehavioral Reviews, page 105744.

[32] Glasser, M. F., Sotiropoulos, S. N., Wilson, J. A., Coalson, T. S., Fischl, B., Andersson, J. L., Xu, J., Jbabdi, S., Webster, M., Polimeni, J. R., et al. (2013). The minimal preprocessing pipelines for the human connectome project. Neuroimage, 80:105–124.

[33] Gonzalez, M. Z., Coppola, A. M., Allen, J. P., and Coan, J. A. (2021). Yielding to social presence as a bioenergetic strategy: Preliminary evidence using fmri. Current Research in Ecological and Social Psychology, 2:100010.

[34] Gordon, E. M., Laumann, T. O., Adeyemo, B., Huckins, J. F., Kelley, W. M., and Petersen, S. E. (2016). Generation and evaluation of a cortical area parcellation from resting-state correlations. Cerebral cortex, 26(1):288–303.

[35] Gordon, E. M., Laumann, T. O., Adeyemo, B., and Petersen, S. E. (2017). Individual variability of the system-level organization of the human brain. Cerebral cortex, 27(1):386–399.

[36] Gross, E. B. and Medina-DeVilliers, S. E. (2020). Cognitive processes unfold in a social context: a review and extension of social baseline theory. Frontiers in Psychology, 11:378.

[37] Guo, T. and Li, X. (2023). Machine learning for predicting phenotype from genotype and environment. Current Opinion in Biotechnology, 79:102853.

[38] Hawkley, L. C. and Cacioppo, J. T. (2007). Aging and loneliness: Downhill quickly? Current Directions in Psychological Science, 16(4):187–191.

[39] Hermans, E. J., Henckens, M. J., Joëls, M., and Fernández, G. (2014). Dynamic adaptation of large-scale brain networks in response to acute stressors. Trends in neurosciences, 37(6):304–314.

[40] Herzberg, M. P., McKenzie, K. J., Hodel, A. S., Hunt, R. H., Mueller, B. A., Gunnar, M. R., and Thomas, K. M. (2021). Accelerated maturation in functional connectivity following early life stress: Circuit specific or broadly distributed? Developmental cognitive neuroscience, 48:100922.

[41] Holmes, C. J., Barton, A. W., MacKillop, J., Galván, A., Owens, M. M., McCormick, M. J., Yu, T., Beach, S. R., Brody, G. H., and Sweet, L. H. (2018). Parenting and salience network connectivity among african americans: a protective pathway for health-risk behaviors. Biological psychiatry, 84(5):365–371.

[42] Holz, N. E., Berhe, O., Sacu, S., Schwarz, E., Tesarz, J., Heim, C. M., and Tost, H. (2022). Early social adversity altered brain functional connectivity and mental health. Biological Psychiatry.

[43] Honegger, K. and de Bivort, B. (2018). Stochasticity, individuality and behavior. Current Biology, 28(1):R8– R12.

[44] House, J. S., Landis, K. R., and Umberson, D. (1988). Social relationships and health. Science, 241(4865):540– 545.

[45] Korgaonkar, M. S., Breukelaar, I. A., Felmingham, K., Williams, L. M., and Bryant, R. A. (2023). Association of neural connectome with early experiences of abuse in adults. JAMA Network Open, 6(1):e2253082–e2253082.

[46] Kuppens, S. and Ceulemans, E. (2019). Parenting styles: A closer look at a well-known concept. Journal of child and family studies, 28(1):168–181.

[47] Li, Y., Liu, Y., Zhao, X., Ren, Y., Hu, W., Yang, Z., and Yang, J. (2024). Static and temporal dynamic in functional connectivity of large-scale brain networks during acute stress regulate stress resilience differently: The promotion role of trait resilience. Neuroscience, 551:132–142.

[48] McLaughlin, K. A., Sheridan, M. A., and Lambert, H. K. (2014). Childhood adversity and neural development: deprivation and threat as distinct dimensions of early experience. Neuroscience & Biobehavioral Reviews, 47:578–591.

[49] Merritt, H., Faskowitz, J., Gonzalez, M. Z., and Betzel, R. F. (2023). Stability of brain-behavior correlation patterns across measures of social support. bioRxiv, pages 2023–03.

[50] Miller, G., Chen, E., and Cole, S. W. (2009). Health psychology: Developing biologically plausible models linking the social world and physical health. Annual review of psychology, 60:501–524.

[51] Moncrieff, J., Cooper, R. E., Stockmann, T., Amendola, S., Hengartner, M. P., and Horowitz, M. A. (2024). Difficult lives explain depression better than broken brains. Molecular Psychiatry, pages 1–4.

[52] Morawetz, C., Berboth, S., and Bode, S. (2021). With a little help from my friends: The effect of social proximity on emotion regulation-related brain activity. Neuroimage, 230:117817.

[53] Mwilambwe-Tshilobo, L., Ge, T., Chong, M., Ferguson, M. A., Misic, B., Burrow, A. L., Leahy, R. M., and Spreng, R. N. (2019). Loneliness and meaning in life are reflected in the intrinsic network architecture of the brain. Social cognitive and affective neuroscience, 14(4):423–433.

[54] Mwilambwe-Tshilobo, L., Setton, R., Bzdok, D., Turner, G. R., and Spreng, R. N. (2022). Age differences in functional brain networks associated with loneliness and empathy. Network Neuroscience, pages 1–60.

[55] Oshri, A., Howard, C. J., Zhang, L., Reck, A., Cui, Z., Liu, S., Duprey, E., Evans, A. I., Azarmehr, R., and Geier, C. F. (2024). Strengthening through adversity: The hormesis model in developmental psychopathology. Development and Psychopathology, pages 1–17.

[56] Pagliaccio, D., Luby, J. L., Bogdan, R., Agrawal, A., Gaffrey, M. S., Belden, A. C., Botteron, K. N., Harms, M. P., and Barch, D. M. (2015). Amygdala functional connectivity, hpa axis genetic variation, and life stress in children and relations to anxiety and emotion regulation. Journal of abnormal psychology, 124(4):817.

[57] Petri, G., Expert, P., Turkheimer, F., Carhart-Harris, R., Nutt, D., Hellyer, P. J., and Vaccarino, F. (2014). Homological scaffolds of brain functional networks. Journal of The Royal Society Interface, 11(101):20140873.

[58] Pfeiffer, U. J., Timmermans, B., Vogeley, K., Frith, C. D., and Schilbach, L. (2013). Towards a neuroscience of social interaction.

[59] Power, J. D., Fair, D. A., Schlaggar, B. L., and Petersen, S. E. (2010). The development of human functional brain networks. Neuron, 67(5):735–748.

[60] Pozzi, E., Vijayakumar, N., Byrne, M. L., Bray, K. O., Seal, M., Richmond, S., Zalesky, A., and Whittle, S. L. (2021). Maternal parenting behavior and functional connectivity development in children: A longitudinal fmri study. Developmental cognitive neuroscience, 48:100946.

[61] Prompiengchai, S. and Dunlop, K. (2024). Breakthroughs and challenges for generating brain network-based biomarkers of treatment response in depression. Neuropsychopharmacology, pages 1–16.

[62] Ragone, E., Tanner, J., Jo, Y., Zamani Esfahlani, F., Faskowitz, J., Pope, M., Coletta, L., Gozzi, A., and Betzel, R. (2024). Modular subgraphs in large-scale connectomes underpin spontaneous co-fluctuation events in mouse and human brains. Communications Biology, 7(1):126.

[63] Rakesh, D., Allen, N. B., and Whittle, S. (2021a). Longitudinal changes in within-salience network functional connectivity mediate the relationship between childhood abuse and neglect, and mental health during adolescence. Psychological medicine, pages 1–13.

[64] Rakesh, D., Seguin, C., Zalesky, A., Cropley, V., and Whittle, S. (2021b). Associations between neighborhood disadvantage, resting-state functional connectivity, and behavior in the adolescent brain cognitive development study: the moderating role of positive family and school environments. Biological Psychiatry: Cognitive Neuroscience and Neuroimaging, 6(9):877–886.

[65] Rakesh, D., Zalesky, A., and Whittle, S. (2021c). Similar but distinct–effects of different socioeconomic indicators on resting state functional connectivity: Findings from the adolescent brain cognitive development (abcd) study®. Developmental cognitive neuroscience, 51:101005.

[66] Rakesh, D., Zalesky, A., and Whittle, S. (2022). The role of school environment in brain structure, connectivity, and mental health in children: A multimodal investigation. Biological Psychiatry: Cognitive Neuroscience and Neuroimaging.

[67] Ramphal, B., DeSerisy, M., Pagliaccio, D., Raffanello, E., Rauh, V., Tau, G., Posner, J., Marsh, R., and Margolis, A. E. (2020a). Associations between amygdala-prefrontal functional connectivity and age depend on neighborhood socioeconomic status. Cerebral Cortex Communications, 1(1):tgaa033.

[68] Ramphal, B., Whalen, D. J., Kenley, J. K., Yu, Q., Smyser, C. D., Rogers, C. E., and Sylvester, C. M. (2020b). Brain connectivity and socioeconomic status at birth and externalizing symptoms at age 2 years. Developmental cognitive neuroscience, 45:100811.

[69] Repetti, R. L., Taylor, S. E., and Seeman, T. E. (2002). Risky families: family social environments and the mental and physical health of offspring. Psychological bulletin, 128(2):330.

[70] Rice, M., Merritt, H., and Gonzalez, M. Z. (2022). The developmental social calibration of reinforcement sensitivity and internalizing symptoms in emerging adulthood. Psychological Science.

[71] Richmond, S., Johnson, K. A., Seal, M. L., Allen, N. B., and Whittle, S. (2016). Development of brain networks and relevance of environmental and genetic factors: a systematic review. Neuroscience & Biobehavioral Reviews, 71:215–239.

[72] Rudolph, K. D., Davis, M. M., Skymba, H. V., Modi, H. H., and Telzer, E. H. (2021). Social experience calibrates neural sensitivity to social feedback during adolescence: A functional connectivity approach. Developmental Cognitive Neuroscience, 47:100903.

[73] Saxbe, D. E., Beckes, L., Stoycos, S. A., and Coan, J. A. (2020). Social allostasis and social allostatic load: A new model for research in social dynamics, stress, and health. Perspectives on Psychological Science, 15(2):469–482.

[74] Schilbach, L., Timmermans, B., Reddy, V., Costall, A., Bente, G., Schlicht, T., and Vogeley, K. (2013). Toward a second-person neuroscience 1. Behavioral and Brain Sciences, 36(4):393–414.

[75] Seeman, T. E. and McEwen, B. S. (1996). Impact of social environment characteristics on neuroendocrine regulation. Psychosomatic medicine, 58(5):459–471.

[76] Sheridan, M. A. and McLaughlin, K. A. (2014). Dimensions of early experience and neural development: deprivation and threat. Trends in cognitive sciences, 18(11):580–585.

[77] Spreng, R. N., Dimas, E., Mwilambwe-Tshilobo, L., Dagher, A., Koellinger, P., Nave, G., Ong, A., Kernbach, J. M., Wiecki, T. V., Ge, T., et al. (2020). The default network of the human brain is associated with perceived social isolation. Nature communications, 11(1):1–11.

[78] Sripada, R. K., Swain, J. E., Evans, G. W., Welsh, R. C., and Liberzon, I. (2014). Childhood poverty and stress reactivity are associated with aberrant functional connectivity in default mode network. Neuropsychopharmacology, 39(9):2244–2251.

[79] Steinberg, L., Blatt-Eisengart, I., and Cauffman, E. (2006). Patterns of competence and adjustment among adolescents from authoritative, authoritarian, indulgent, and neglectful homes: A replication in a sample of serious juvenile offenders. Journal of research on adolescence, 16(1):47–58.

[80] Sydnor, V. J., Larsen, B., Seidlitz, J., Adebimpe, A., Alexander-Bloch, A. F., Bassett, D. S., Bertolero, M. A., Cieslak, M., Covitz, S., Fan, Y., et al. (2023). Intrinsic activity development unfolds along a sensorimotor–association cortical axis in youth. Nature Neuroscience, pages 1–12.

[81] Thomason, M. E., Marusak, H. A., Tocco, M. A., Vila, A. M., McGarragle, O., and Rosenberg, D. R. (2015). Altered amygdala connectivity in urban youth exposed to trauma. Social cognitive and affective neuroscience, 10(11):1460–1468.

[82] Tian, Y. E., Di Biase, M. A., Mosley, P. E., Lupton, M. K., Xia, Y., Fripp, J., Breakspear, M., Cropley, V., and Zalesky, A. (2023). Evaluation of brain-body health in individuals with common neuropsychiatric disorders. JAMA psychiatry.

[83] Tottenham, N. (2014). The importance of early experiences for neuro-affective development. The neurobiology of childhood, pages 109–129.

[84] Uchino, B. N., Cacioppo, J. T., and Kiecolt-Glaser, J. K. (1996). The relationship between social support and physiological processes: a review with emphasis on underlying mechanisms and implications for health. Psychological bulletin, 119(3):488.

[85] Uchino, B. N., Landvatter, J., Zee, K., and Bolger, N. (2020). Social support and antibody responses to vaccination: a meta-analysis. Annals of Behavioral Medicine, 54(8):567–574.

[86] Uchino, B. N., Trettevik, R., Kent de Grey, R. G., Cronan, S., Hogan, J., and Baucom, B. R. (2018). Social support, social integration, and inflammatory cytokines: A meta-analysis. Health Psychology, 37(5):462.

[87] Van Essen, D. C., Smith, S. M., Barch, D. M., Behrens, T. E., Yacoub, E., Ugurbil, K., Consortium, W.-M. H., et al. (2013). The wu-minn human connectome project: an overview. Neuroimage, 80:62–79.

[88] Vedechkina, M., Astle, D. E., and Holmes, J. (2024). Dimensions of early life adversity and their associations with functional brain organisation. Imaging Neuroscience.

[89] Wang, F., Gao, Y., Han, Z., Yu, Y., Long, Z., Jiang, X., Wu, Y., Pei, B., Cao, Y., Ye, J., et al. (2023). A systematic review and meta-analysis of 90 cohort studies of social isolation, loneliness and mortality. Nature Human Behaviour, pages 1–13.

[90] Webb, E. K., Cardenas-Iniguez, C., and Douglas, R. (2022). Radically reframing studies on neurobiology and socioeconomic circumstances: A call for social justice-oriented neuroscience. Frontiers in Integrative Neuroscience, 16.

[91] Wen, J., Tian, Y. E., Skampardoni, I., Yang, Z., Cui, Y., Anagnostakis, F., Mamourian, E., Zhao, B., Toga, A. W., Zalesky, A., et al. (2024). The genetic architecture of biological age in nine human organ systems. Nature Aging, pages 1–18.

[92] Williams, W. C., Morelli, S. A., Ong, D. C., and Zaki, J. (2018). Interpersonal emotion regulation: Implications for affiliation, perceived support, relationships, and well-being. Journal of personality and social psychology, 115(2):224.

[93] Wymbs, N. F., Orr, C., Albaugh, M. D., Althoff, R. R., O’Loughlin, K., Holbrook, H., Garavan, H., Montalvo-Ortiz, J. L., Mostofsky, S., Hudziak, J., et al. (2020). Social supports moderate the effects of child adversity on neural correlates of threat processing. Child abuse & neglect, 102:104413.

[94] Xiao, M., Chen, X., Yi, H., Luo, Y., Yan, Q., Feng, T., He, Q., Lei, X., Qiu, J., and Chen, H. (2021). Stronger functional network connectivity and social support buffer against negative affect during the covid-19 outbreak and after the pandemic peak. Neurobiology of stress, 15:100418.

[95] Yang, Z., Zuo, X.-N., McMahon, K. L., Craddock, R. C., Kelly, C., de Zubicaray, G. I., Hickie, I., Bandettini, P. A., Castellanos, F. X., Milham, M. P., et al. (2016). Genetic and environmental contributions to functional connectivity architecture of the human brain. Cerebral cortex, 26(5):2341–2352.

[96] Young, E. S., Frankenhuis, W. E., and Ellis, B. J. (2020). Theory and measurement of environmental unpredictability. Evolution and Human Behavior, 41(6):550–556.

[97] Zaki, J. and Williams, W. C. (2013). Interpersonal emotion regulation. Emotion, 13(5):803.

